# DUCs are C2 domain containing plant-specific deubiquitinases stabilizing endocytic cargo at the plasma membrane

**DOI:** 10.64898/2025.12.04.692294

**Authors:** Vanessa Lachmayer, Karin Vogel, Meleen Scholte-Wassink, Thomas Hermanns, Tobias Bläske, Jan M. Gebauer, Ulrich Baumann, Erika Isono, Kay Hofmann

**Author notes:** shared first authors (Vanessa Lachmayer, Karin Vogel). corresponding authors Correspondence should be addressed to Kay Hofmann and Erika Isono.

## Abstract

Deubiquitinases (DUBs) remove ubiquitin modifications from proteins in a substrate- or linkage-selective manner and regulate numerous cell-biological processes, including endocytosis. By performing homology-agnostic bioinformatical screens for undescribed deubiquitinase classes, we identified the plant-specific DUC (<u>d</u>e<u>u</u>biquitinase with <u>C</u>2) family, whose members are highly selective for cleaving K63-linked ubiquitin chains. The crystal structure of the Arabidopsis member AtDUC1 reveals that DUC enzymes display a papain-like fold with a characteristic DUB-like active site, despite the apparent absence of sequence similarity to other deubiquitinase families. The K63 linkage specificity is attributed to the recognition of both distal and proximal ubiquitin units via highly conserved contact residues. Arabidopsis DUC members localize to the plasma membrane by virtue of their lipid-binding C2-like domains. In protoplast experiments, all three Arabidopsis DUC enzymes demonstrate the capacity to stabilize an endocytotic model cargo at the plasma membrane, a process that requires both the C2-mediated membrane association and catalytic DUB activity.

## Introduction

Protein ubiquitination is a post-translational modification that regulates the stability, localization, and other properties of the modified proteins. The ubiquitin modifier itself is subject to further ubiquitination on seven different lysine side chains and the N-terminal amino group, resulting in ubiquitin chains of different linkage types, which influence the fate of the substrate in different ways ^1^. The highly-abundant K48-linked ubiquitin chains target nuclear and cytoplasmic substrates for proteasomal degradation. By contrast, K63-linked chains have different functions in nuclear and cytoplasmic protein pools: In the nucleus, K63 chains regulate the localization of proteins to DNA damage sites. On the other hand, K63-linked chains attached to the cytoplasmic portion of membrane receptors induce their endocytosis and are crucially involved in the decision between receptor recycling and lysosomal degradation ^2,3^. Other chain types are involved in regulating other processes, including the cell cycle (K11), mitophagy (K6), and immunity (M1). A large number of ubiquitin ligases (E3s) and deubiquitinases (DUBs), which are enzymes responsible for installing/removing ubiquitin-based signals, are found in all eukaryotes. These enzymes can be specific for the modified substrate and/or for the linkage type of the modifying ubiquitin chain.

Seven deubiquitinase classes are found throughout all eukaryotic kingdoms, among them the JAMM-type metalloproteases and six types of cysteine proteases that go by the names USP, UCH, OTU, MJD/Josephin, MINDY and ZUFSP/ZUP ^4^. Among these enzyme families, the OTU-type deubiquitinases are known for their linkage-specificity ^5^. Recently, an eighth eukaryotic DUB class (VTD) was identified, which has an OTU-like propensity for linkage specificity but is only found in selected eukaryotic taxa and is absent from mammals and plants ^6^. Individual members of other DUB classes can also be linkage-specific, as exemplified by mammalian AMSH, a highly K63-selective metallo-DUB ^7^, which regulates the fate of endocytosed cargo at the endosome level ^8^. In plants, endocytosis is also regulated by K63-linked ubiquitin chains ^9^, although there are major mechanistic differences. ESCRT-0, the primary recognition complex for ubiquitinated endocytosis cargo in animals and fungi, is missing in plants and is functionally replaced by other ubiquitin-binding proteins ^10^. In contrast, the other ESCRT complexes, which guide ubiquitinated cargo to the endosome, are conserved in plants, and the deubiquitinase AMSH3 is associated with endosomes and appears to function similarly to its metazoan counterpart ^11^. Four additional deubiquitinases have been implicated in the regulation of plant endocytosis: cytoplasmic UBP12 and UBP13 and plasma membrane-associated OTU11 and OTU12 ^11,12^. However, they show no selectivity for K63-linked chains, and their exact mode of action is insufficiently understood.

A recent comprehensive analysis of plant deubiquitinases ^13^ revealed broad similarities to the situation in animals and other eukaryotic taxa ^4^. Although the DUB complement of Arabidopsis thaliana is somewhat smaller than that of humans, the two genomes contain identical DUB classes in similar proportions. The identification of non-canonical DUB classes in bacteria ^14,15^ and the taxonomically restricted VTD family in insects, nematodes, and fungi, but not in mammals or plants ^6^, suggests that there may be new species-restricted DUB classes that remain to be discovered. Previous efforts to identify DUB families have largely relied on activity-based probes ^16–18^ or homology-based bioinformatical screens ^6,19–21^. However, the former approach usually misses linkage-specific DUBs, whereas the latter is unlikely to discover fundamentally new DUB architectures. To circumvent these restrictions, we performed a homology-agnostic bioinformatics screen based on the structural co-modelling of candidate proteins with ubiquitin and analyzed the predicted binding geometry.

When this method was applied to the proteome of *Arabidopsis thaliana*, we recovered most of the established cysteine DUBs and several uncharacterized members of established DUB families, as well as an uncharacterized Arabidopsis ORF with confidently predicted DUB-like properties. Subsequently, this singular finding could be expanded to a family of three Arabidopsis DUB candidates and numerous homologues from other plants, unrelated by sequence to any known DUB family. The experimental validation of representative enzymes from *Arabidopsis* and *Physcomitrium* found a strong K63-directed deubiquitinase activity. Structure-based mutagenesis of key residues revealed the determinants of DUB activity and K63-specificity. N-terminal of their enzymatic domain, all members of this plant-specific deubiquitinase family contain a C2 domain, which is responsible for recruitment of the enzymes to the plasma membrane, where the deubiquitinase activity is required to stabilize endocytotic cargo.

## Results

### Bioinformatical discovery of the DUC deubiquitinases

Since previously employed bioinformatical screens for new deubiquitinases were based on sequence homology and thus unable to discover fundamentally new DUB families, we set up a homology-independent screen based on molecular modeling ^22^. While using Alphafold-2 ^23^ for modelling members of various DUB classes with ubiquitin, we had observed that the ubiquitin is consistently placed at the S1 recognition site, with its C-terminus adjacent to the catalytic cysteine – at least in cases where the catalytic domain is amenable to Alphafold modeling. To screen the protein-coding part of the *Arabidopsis thaliana* genome for potential new cysteine-type deubiquitinases, we set up a multi-step prediction pipeline that minimizes the number of necessary co-models. In the first step, the *A. thaliana* subset of the AlphaFold Protein Structure Database ^24^ was used, which contains 27,434 single-protein structures (Supplementary Data File 1, Sheet 1). These structures were screened for residue clusters that might constitute the active sites of cysteine proteases, consisting of spatially close cysteine, histidine, and aspartate/asparagine residues. Overall, 2291 such clusters with inter-residue spacing < 4.2 Å were obtained (Supplementary Data File 1, Sheet 2), of which only a minority is expected to represent genuine cysteine proteases. To further enrich this list for deubiquitinases, we focused on active site candidates harboring the DUB-typical gatekeeper motif ^21^, by requiring that the predicted active site histidine residue is directly followed by an aromatic residue. After this filtering step, 152 proteins were retained, among which the vast majority were known Arabidopsis deubiquitinases ^13^. These proteins (known DUBs and new candidates) were subjected to co-modeling with ubiquitin, and the resulting models were evaluated using AlphaFold’s performance metric ipTM and the position of the modelled ubiquitin C-terminus relative to the catalytic cysteine. The results are summarized in Supplementary Data File 1, Sheet 3.

Of the 152 modelled proteins (Supplementary Data File 2), 58 had ipTM scores greater than 0.75, suggesting that they could form a complex with ubiquitin. In 54 of the calculated models, the ubiquitin C-terminus was found within 4 Å of the predicted active-site thiol group. 48 proteins fulfilled both criteria, the majority of which are known proteases for ubiquitin and/or ubiquitin-like modifiers ^13^. A single uncharacterized protein (At2g25460) was found among the high-confidence predictions, with an ipTM value of 0.96 and a distance of 3.3 Å between the ubiquitin C-terminus and Cys-318 of the DUB candidate. Sequence analysis of At2g25460 showed that it is a member of a large plant-specific protein family, with two additional genes in A. thaliana (At3g11760 and At5g04860) that were not identified on the screen because they were missing from the downloadable dataset of *Arabidopsis* models. Multiple members of the gene family are found in all major taxa of land plants, including dicotyledon and monocotyledon angiosperms, gymnosperms, and bryophytes; single copies are found in selected algae (Fig 1a). A common feature of all embryophyte-derived members is the presence of a C2-like domain at the N-terminus, followed by a flexible linker, and a predicted DUB domain at the C-terminus (Fig 1b). In some algal members, the C2 domain is either highly divergent or absent. Based on this conserved architecture, we named this new family DUC (<u>DU</u>Bs with <u>C</u>2 domain) and refer to the *A. thaliana* members as AtDUC1 (At2g25460), AtDUC2 (At3g11760), and AtDUC3 (At5g04860). In addition to the three *Arabidopsis* proteins, we selected the evolutionarily distant *Physcomitrium* protein PpDUC (PHYPA_013469) for further characterization.

**Figure 1:**
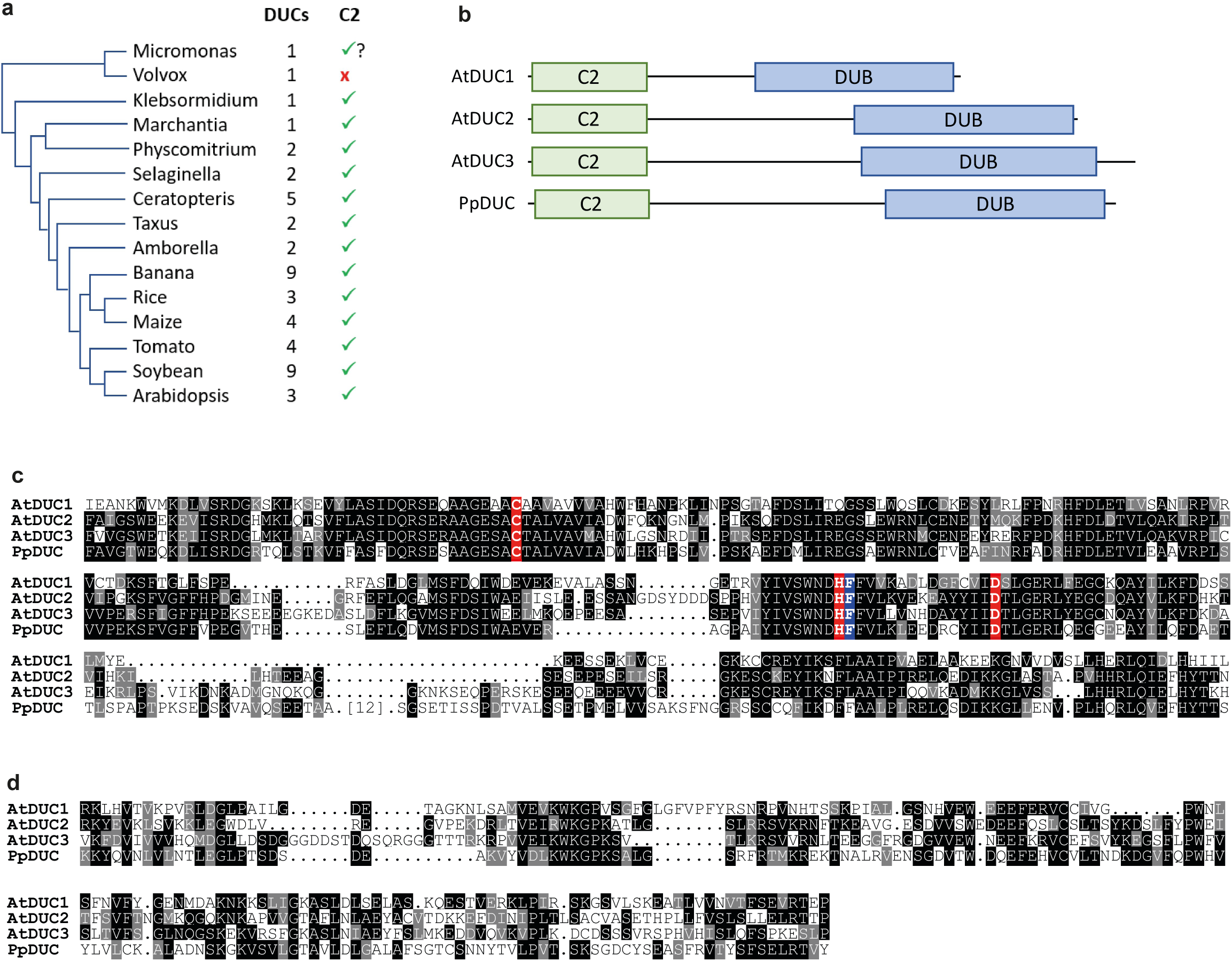
Bioinformatical characterization of the plant DUC family. **(a)** Dendrogram of species representing different plant clades and families. The number of DUC family members and the presence of an N-terminal C2 domain are indicated. **(b)** Domain scheme of the four DUC members studied here. The approximate positions and sizes of the C2 (lipid-binding) and DUB (deubiquitinase) domains are indicated. **(c)** Sequence alignment of the catalytic deubiquitinase domains. Invariant and conservatively replaced residues are shown on a black or gray background, respectively. The catalytic triad is highlighted in red and the gatekeeper residue in blue. (d) Sequence alignment of the C2-like domain. The color rendering was performed as described above.

As shown in Fig 1c, the predicted catalytic domain spans about 300 residues and is remarkably well conserved with 47-60% residue identity between the *Arabidopsis* and *Physcomitrium*-derived proteins. The predicted active site has a Cys-His-Asp arrangement, which is typical of papain-fold proteases. The presence of a conserved aromatic residue (Phe) following the catalytic histidine is a hallmark of DUBs and GlyGly-directed proteases ^21^. No meaningful sequence similarity to other DUBs or proteases was detected, suggesting that the DUC proteins do not belong to any of the established DUB classes. A second region of sequence conservation was found in the N-terminus and corresponds to the predicted C2-like domain (Fig 1d). It is less well conserved (29-35% residue identity) than the catalytic domain, but shows a significant sequence relationship (p<10^-7^) to established C2 domains ^25^, the most similar ones being the N-terminal C2 domain of human EHBP1 ^26^ and other members of the INTERPRO family IPR019448 ^27^.

On the transcript level, At*DUC1*, At*DUC2*, and At*DUC3* are broadly expressed in Arabidopsis tissues ^28^ (Supplemental Fig 2a and b). There are slight differences in the expression pattern depending on tissues. DUC3 is, for example, more strongly expressed in the root and seeds than AtDUC1 and AtDUC2.

### Biochemical characterization of catalytic deubiquitinase domain

To determine the enzymatic activity of AtDUC1-3, their catalytic domain boundaries were identified based on sequence alignments (Fig 1c) and their models from the AlphaFold Protein Structure Database ^24^. The resulting catalytic constructs, AtDUC1^277–546^, AtDUC2^412-702^, and AtDUC3^427-733^, were expressed as Smt3 fusion proteins in E. coli, purified, and processed to yield catalytically active protein fragments, which were subjected to a number of enzymatic assays (Fig 2). Most deubiquitinases react with the activity-based probe Ub-PA ^29^, resulting in a covalent adduct between DUB and ubiquitin. As shown in Fig 2a-c, the catalytic domains of all three AtDUCs reacted readily with Ub-PA, as visualized by an upward shift of the DUB band by ~8kDa. When testing reactivity with analogous probes based on ubiquitin-like modifiers, a weak reaction with Nedd8-PA, but no reactivity towards ISG15- or SUMO-derived probes was detected (Fig 2a-c). To assess chain cleavage and evaluate a potential linkage specificity, purified catalytic fragments of AtDUCs were incubated with a diubiquitin panel of all eight chain types. AtDUC1^277-546^ efficiently cleaved K63-linked di-ubiquitin to near-completion within 10 min (Fig 2d). By contrast, other linkage types were hardly cleaved, as no reduction of those di-ubiquitin species was observed. The paralogs AtDUC2^412-702^ and AtDUC3^427-733^ showed a similar K63-preference, albeit with some trace reactivity against K11-linked di-ubiquitin (Fig 2e-f).

**Figure 2.**
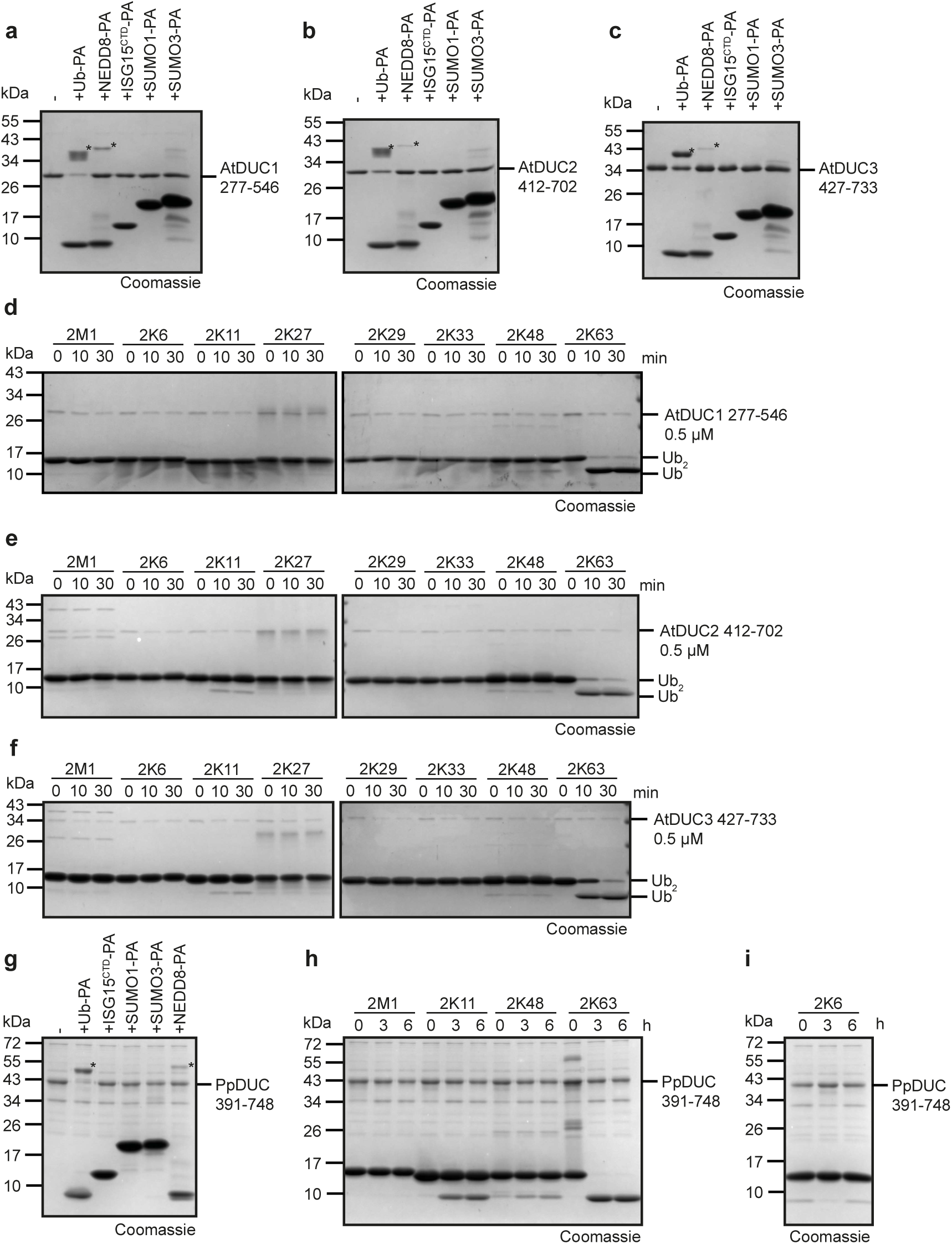
Deubiquitinase activity of plant DUCs. **(a,b,c)** Catalytic fragments of AtDUC1^277-546^, AtDUC2^412-702^, and AtDUC3^423-733^ were incubated overnight with activity-based probes for ubiquitin (Ub-PA), NEDD8 (NEDD8-PA), the C-terminal domain of ISG15 (ISG15^CTD^-PA), and two SUMO forms (SUMO1-PA, SUMO3-PA). Size shifts caused by reactivity are indicated by an asterisk (*). **(d,e,f)** Catalytic fragments ofAtDUC1, AtDUC2, and AtDUC3 were incubated for the indicated time periods with di-ubiquitin species of the indicated linkage types. Reactivity was visualized by the disappearance of di-ubiquitin (Ub_2_) and appearance of mono-ubiquitin (Ub) bands in Coomassie-stained gels. **(g)** Activity-based probe assay for Physcomitrium catalytic fragment PpDUC^391-748^. The probes and lettering were used as in panels a-c. **(h,i)** Catalytic fragment PpDUC^391-748^ was incubated for 6h with di-ubiquitin species of M1, K11, K48, K63, and K6 linkages. Lettering as in panels (d) and (e).

To address the question if K63-selectivity is a hallmark of the DUC family, we also examined an evolutionarily distant member from the bryophyte *Physcomitrium patens*. The catalytic fragment PpDUC^391-748^ showed similar reactivity towards activity-based probes as the *Arabidopsis* proteins, with strong reactivity against Ub-PA, weak reactivity against Nedd8-PA, and no reactivity towards ISG15-or SUMO-derived probes (Fig 2g). The cleavage of differently linked di-ubiquitin species required longer incubation times than that of AtDUCs, but revealed a similar specificity profile with strong reactivity against K63-linked chains and trace reactivity against K11- and K48-linkages (Fig 2h,i). In summary, our experimental data suggest that members of the plant DUC family are active deubiquitinases with a strong preference for cleaving K63-linkages.

### A structural basis for catalysis

The absence of sequence similarity between DUC members and other peptidases and isopeptidases suggests that DUCs form a novel deubiquitinase class unrelated to the seven established cysteine DUB classes. However, all cysteine DUBs and proteases for ubiquitin-like modifiers have a papain fold, indicating a distant evolutionary relationship that has diverged beyond being recognizable at the sequence level. To study the evolutionary origins of the DUC family, we attempted to determine the crystal structure of a representative DUC-type enzyme. To date, all attempts to crystallize DUCs in complex with a di-ubiquitin substrate or as a covalent adduct with ubiquitin-PA have failed, but crystals of the isolated AtDUC1^277-546^ catalytic fragment diffracted well and allowed to solve a structure at approximately 2.0 Å resolution (Fig 3a). Three internal loops (309-316, 356-372, 462-464) and the last two residues were not clearly resolved and thus not built. The remaining 240 residues, however, show a papain-like fold with little deviation from the Alphafold-2 model used for its discovery (0.84 Å RMSD over the entire sequence, Supplementary Fig 1). The active site residues Cys-318, His-441, and Asp-456 have an arrangement typical of papain-fold deubiquitinases and show the expected degree of sequence conservation (Fig 1c). Mutation of any active site residue to alanine (C318A, H441A, and D456A) led to a complete loss of catalytic activity against K63-linked diubiquitin, while mutation of the gatekeeper residue (F442A) led to strongly reduced activity (Fig 3b).

**Figure 3.**
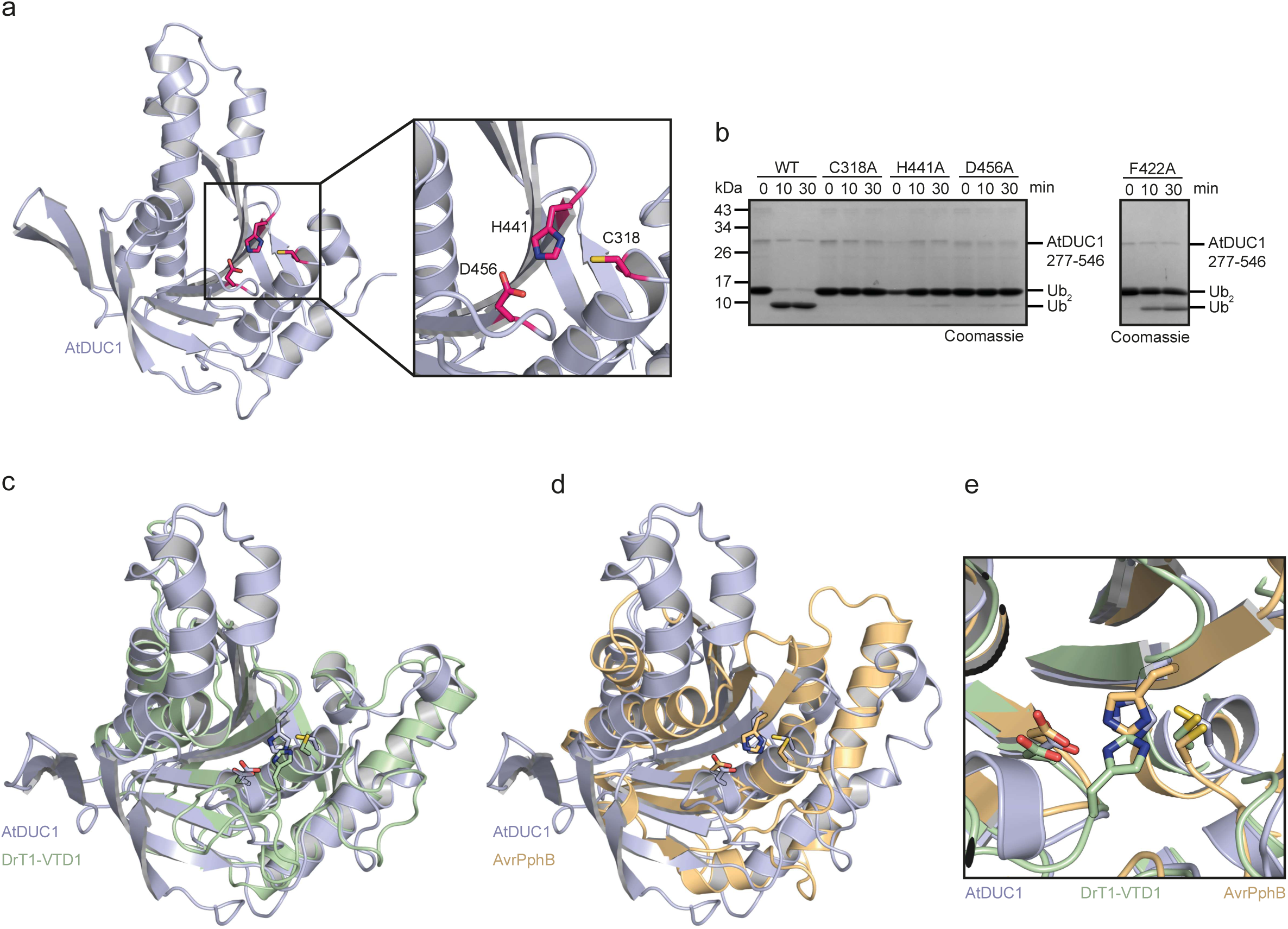
Crystal structure of AtDUC1. **(a)** Structure of the catalytic domain AtDUC1^277-546^ shown in cartoon representation with the catalytic triad residues highlighted as red sticks. **(b)** Wild-type catalytic fragment AtDUC1^277-546^ and variants with individual catalytic residues mutated to alanine were incubated with K63-linked di-ubiquitin and analyzed by Coomassie-stained SDS-PAGE. **(c)** Comparison of the AtDUC1 structure (light purple) with that of DrT1-VTD1 (light green); catalytic residues are shown as sticks, and the remaining structure is shown as a cartoon. **(d)** Comparison of the AtDUC1 structure (light purple) with that of AvrPphB (dark yellow). Catalytic residues are shown as sticks, and the remaining structure is shown as a cartoon. **(e)** Zoom-in on the superimposed catalytic center of AtDUC1 (light purple), DrT1-VTD1 (light green), and AvrPphB (dark yellow).

Structural database searches using the DALI software ^30^ showed significant similarity to two different protease families. The best-scoring match was DrT1VTD1 (PDB:8ADD), a deubiquitinase encoded by a zebrafish transposon ^6^. The Z-score for structural similarity was 17.5, indicating high significance, whereas the associated sequence identity of 13% was remarkably low. The following matches with Z-scores between 14.5 and 16.0 were other members of the VTD family. Additional significant Z-scores in the range of 7.0 to 10.5 were found for the bacterial avirulence protein AvrPphB (PDB:1UKF) ^31^ and other members of the YopT-like peptidase family ^32^. The structural superposition of AtDUC1 and DrT1VTD1 with an RMSD of 2.4 Å over 186 residues showed a convincing resemblance within the papain-fold core and adjacent structural elements (Fig 3c), but revealed major differences in the active site of the two proteins. Like all other VTD-type DUBs, DrT1VTD1 has a Cys-Asp-His catalytic triad, where the catalytic histidine is contributed by the same β-strand as the catalytic aspartate and lacks an aromatic gatekeeper motif ^6^. By contrast, the Cys-His-Asp active site of AtDUC1 resembles that of papain and most DUB classes. The structure of AvrPphB, which has no deubiquitinase activity, can also be convincingly superimposed on the AtDUC1 structure (RMSD of 3.8 Å over 171 residues, Fig 3d) and shows the same active site arrangement, albeit without the gatekeeper motif. In summary, the plant DUC family, the VTD family, and the bacterial YopT family are structurally related but employ different active site architectures (Fig 3e). Owing to the lack of sequence similarity, their evolutionary relationships are difficult to establish.

### Structural Basis of K63-specificity

Because no structures of DUC enzymes in complex with their substrates are available, we generated structural models of AtDUC1 with two unconnected ubiquitin molecules using Alphafold-3 ^33^. The resulting model showed both ubiquitin units bound with high confidence to the S1 and S1’ sites of the deubiquitinase, with the Lys-63 ε-amino group of the proximal (S1’) ubiquitin adjacent to the C-terminus of the distal (S1) ubiquitin and the active site cysteine of AtDUC1 (Fig 4a, Supplementary Data File 3). The plausibility of this model is supported by the high confidence of the two ubiquitin interfaces (ipTM values of 0.90 and 0.87 for distal and proximal ubiquitin, respectively) and the concordance with the experimentally determined K63-selectivity of the enzyme. For experimental validation of the predicted substrate recognition mode, a covalent K63-linkage was artificially introduced, and the resulting strain was relaxed by energy minimization. Based on this refined model, residues at the S1 and S1’ interfaces were selected, mutagenized, and tested for their effects on catalytic activity.

**Figure 4.**
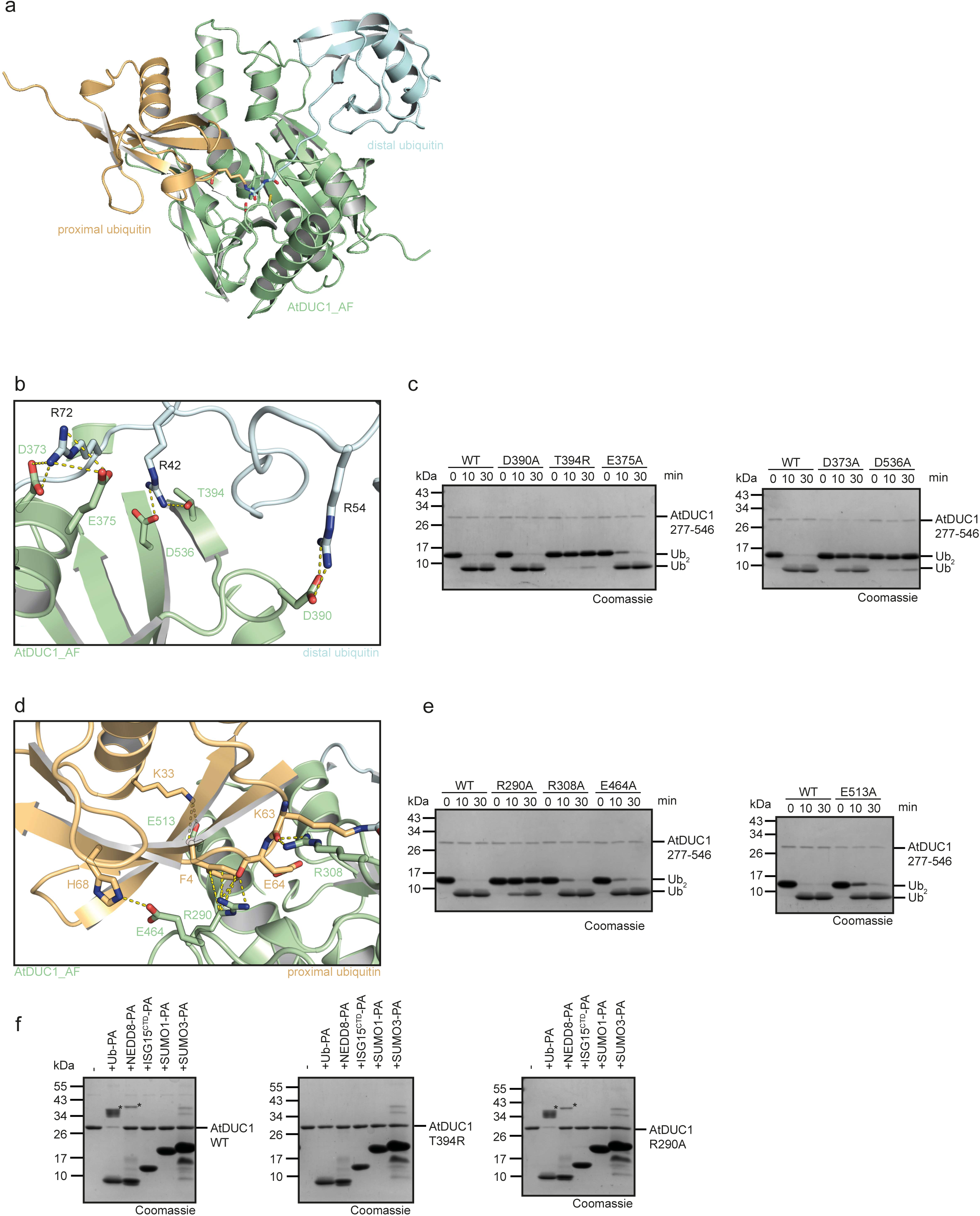
Structural basis of K63 recognition. **(a)** Cartoon representation of the Alphafold-3 model of AtDUC1 (green) in complex with S1-bound distal ubiquitin (light blue) and S1’-bound proximal ubiquitin (dark yellow). The K63 isopeptide linkage and catalytic residues are shown in stick representation. **(b)** Cartoon representation of the S1 binding interface. Residues selected for mutagenesis and their interaction partners are shown as sticks and are labeled. **(c)** Enzymatic activity assays (cleavage of K63-linked di-ubiquitin) for AtDUC1 residues involved in S1 recognition. **(d)** cartoon representation of the S1’ binding interface. Residues selected for mutagenesis and their interaction partners are shown as sticks and are labeled. **(e)** Enzymatic activity assays (cleavage of K63-linked di-ubiquitin) for AtDUC1 residues involved in S1’ recognition. **(f)** Incubation of wildtype AtDUC1^277-546^, S1-interface mutant T394A, and S1’-interface mutant R290A with activity-based probes for ubiquitin (Ub-PA) and ubiquitin-related modifiers. Upshifted bands indicate reactivity and are labeled with asterisks (*).

The S1 interface for the distal ubiquitin unit is characterized by salt bridges and hydrogen bonds between the positively charged ubiquitin residues Arg-42, Arg-54, and Arg-72 and the adjacent acidic AtDUC1 residues Asp-373, Glu-375, Asp-390, and Asp-535 (Fig 4b). The latter four residues were replaced with alanine and tested for activity. In addition, the disruptive variant T394R was generated, as Thr-394 interacts with Arg-42 of ubiquitin, and the mutated version is expected to be incompatible with ubiquitin binding at the S1 position. When testing the enzyme variants for cleavage of K63-linked di-ubiquitin, D390A was as active as the wild type, and E375A showed only a modest effect. In contrast, D373A showed severely reduced activity, whereas D536A and the disruptive T394R variants were mostly inactive (Fig 4c).

In the S1’ interface for the proximal ubiquitin, Arg-290 forms a salt bridge with Glu-64 of ubiquitin (Fig 4d), whose importance for substrate recognition is underscored by the barely active R290A mutant (Fig 4e). Additional contacts between AtDUC1 Arg308 and the ubiquitin main chain near Lys-63, between Glu-513 and ubiquitin Lys-33, or between Glu-464 and ubiquitin Lys-6 (Fig 4d) appear to be less crucial, as the individual point mutants R308A, E315A, and E464A showed only moderately reduced activity (Fig 4e). These results show that mutations in the S1’ interface do impede, and in the case of R290A, mostly prevent K63 di-ubiquitin cleavage, highlighting the importance of the K63-specific interaction mode. The different effects of S1 and S1’-recognition are demonstrated by reacting the S1-disruptive T394R variant and the most severe S1’-recognition mutant R290A with the activity-based probe Ub-PA (Fig 4f). This highly reactive mono-ubiquitin probe utilizes only the S1-surface and thus is completely inactive with the T394R variant. As expected, the reactivity against the S1’-deficient R290A variant was not affected.

### C2 domains bind PIP-lipids and confer membrane association

C2 domains are conserved protein domains that bind to phospholipids, often in a Ca²⁺-dependent manner. As the C2 domain can define the cellular localization of proteins, we examined the C2 domain of *Arabidopsis* DUC1, DUC2, and DUC3 for phospholipid binding. When recombinant C2 domains of DUC1, DUC2, and DUC3 were incubated with membranes spotted with different phospholipids in the presence of CaCl_2_, all showed binding to phosphatidylinositol (PI) and to a lesser extent also to phosphatidic acid (PA) (Fig. 5a and Supplementary Fig. 3a). For further experiments, we selected DUC1 and DUC3. We next tested whether the interaction depends on the presence of Ca^2+^ and incubated the C2 domains of DUC1 and DUC3 with 10 µM CaCl_2_ or with 5 mM EGTA. GST-DUC1(C2) bound to PIs and PA also in the presence of EGTA (Fig. 5b), indicating that DUC1 C2 domain can bind membrane lipids also in the absence of Ca^2+^. GST-DUC3(C2) also bound to PIs in a Ca^2+^-independent manner (Supplementary Fig. 3b).

**Figure 5:**
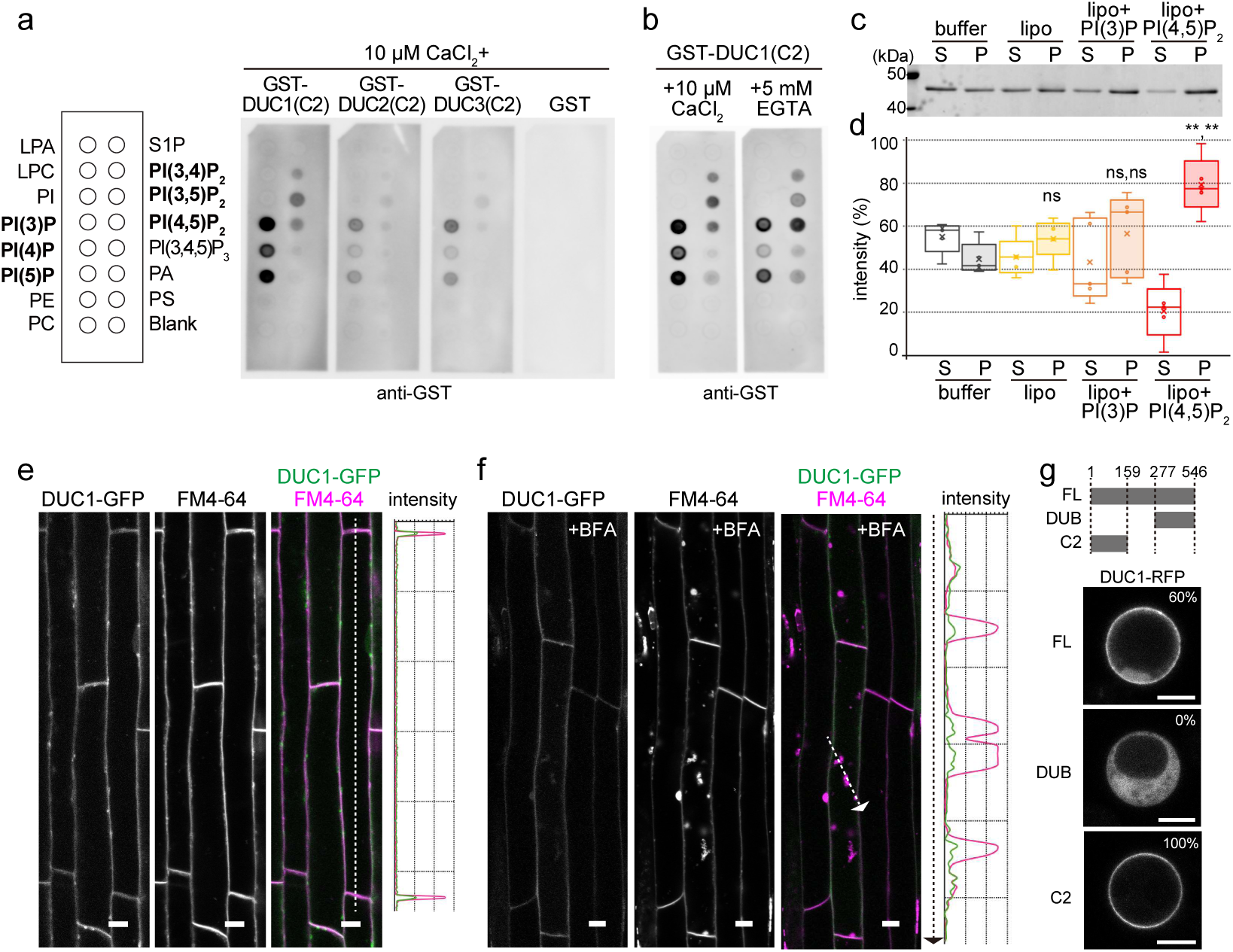
DUCs bind phospholipids through their C2 domain. **(a)** Lipid overlay assays with GST-DUC1(C2), GST-DUC2(C2), GST-DUC3(C2), and GST. GST-fused proteins were incubated with the lipid-spotted membrane in the presence of 10 µM CaCl_2_ and bound proteins were detected with an anti-GST antibody. Lipid species are indicated on the left and thouse bound by GST-DUC1(C2) are indicated in bold. LPA: Lysophosphatidic acid, LPC: Lysophosphatidylcholine, PI: Phosphatidylinositol, PE: Phosphatidylethanolamine, PC: Phosphatidylcholine, S1P: Sphingosine 1-P, PA: Phosphatidic acid, PS: Phosphatidylserine. The experiment was conducted three times and one representative result is shown. **(b)** Lipid overlay assay of GST-DUC1(C2) with either 10 µM CaCl_2_ or 5 mM EGTA and detected as in **(a)**. **(c)** Liposome sedimentation assay with GST-DUC1(C2). GST-DUC1(C2) was incubated with the liposome buffer alone, with liposomes (PC/PE) with or without 5% of PI(3)P or PI(4,5)P_2_ and centrifuged to obtain the supernatant (S) and pellet (P) fractions. SDS-PAGE gels were stained with TCE. The experiment was performed five times and one representative result is shown. **(d)** Quantification of the result in (c). The signal intensity in the supernatant and pellet fractions in relation to the total signal intensity (S+P) is shown. lipo: liposome. Box plot shows the results of five independent experiments. Center line, median; box limits, first and third quartiles; whiskers, 1.5x interquartile range; points, outliers. Comparison of the pellet fraction: lipo/buffer *p*=0.113 (ns: 0.05<*p*), lipo+PI(3)P/buffer *p*=0.254, lipo+PI(3)P/lipo *p*=0.804 (ns: 0.05<*p*), lipo+PI(4,5)P_2_/buffer *p*=0.00179, lipo+PI(4,5)P_2_/lipo *p*=0.00916 (**: 0.001<*p*<0.01). Two-tailed *t*-tests with no equal variance. **(e)** Confocal image of Arabidopsis root epidermis cells expressing *DUC1pro:DUC1-GFP* and incubated with FM4-64 for 5 minutes. The confocal analysis was repeated twice. The signal intensity profile along the dotted line in the merged image is displayed to the right of the panels. Scale bars: 10 µm. **(f)** A representative confocal image of root epidermis cells of Arabidopsis seedling with *DUC1pro:DUC1-GFP* incubated with FM4-64 and brefeldin A (BFA) for 60 minutes before imaging. The analysis was performed twice with five seedlings each. The signal intensity profile along the dotted line in the merged image is shown on the right of the panels. Scale bars: 10 µm. **(g)** The C2 domain of DUC1 is important for the PM localization of DUC1 *in cellula. Arabidopsis* root cell culture-derived protoplasts were transformed with *35Spro:DUC1(FL)-RFP* (*n*=25 cells), *35Spro:DUC1(DUB)-RFP* (*n*=25 cells), or *35Spro:DUC1(C2)-RFP* (*n*=22 cells) and localisation of the RFP-fusion constructs was analysed. Percentage of cells with RFP signals at the PM is indicated. Scale bars: 10 µm. Three independent experiments were conducted, and the result of one representative result is shown.

We subsequently investigated whether DUC1 C2 domain can bind to lipids in the context of lipid bilayers. To this end, we generated liposomes with phosphatidylcholine (PC) and phosphatidylethanolamine (PE) containing no PIPs or 5% of one either PI(3)P, or PI(4,5)P_2_. Recombinant DUC1 (C2) or DUC3 (C2) were then incubated with liposomes and after centrifugation, liposome association was analysed. Both DUC1(C2) and DUC3(C2) did not show a significant increase in the bound fraction when PI(3)P was added to the liposome, but bound PI(4,5)P_2_-containing liposomes significantly more than liposomes without PIPs (Fig. 5c and d, Supplementary Fig. 3c and d). While PI(3)P is found on endosomes and autophagosomes, PI(4,5)P_2_ was shown to be enriched in the plasma membrane of Arabidopsis ^34^, suggesting that DUC1 and DUC3 might localize to PI(4,5)P_2_-containing cellular membranes.

To investigate the localization of DUC1 and DUC3, we generated native promoter-driven GFP fusion constructs to establish stable Arabidopsis lines *DUC1pro:DUC1-GFP* and *DUC3pro:DUC3-GFP*. When the localization of DUC1 and DUC3 was studied in root epidermis cells of Arabidopsis seedlings, DUC1 and DUC3 localization was observed at the plasma membrane (Fig 5e and Supplementary Fig 3e), which was corroborated for DUC1 by co-staining with the lipophilic dye FM4-64 upon 1 minute of incubation prior to imaging (Fig 5e). To test the possibility that DUC1 also localizes to endosomes, we treated DUC1-GFP expressing seedlings with 50 µM brefeldin A (BFA) for 1 hour. BFA is a mycotoxin that inhibits ARF-GEF functions and causes accumulation of early endosome/TGN components to accumulate in a so-called BFA body. While FM4-64 accumulated in BFA bodies, DUC1-GFP signals did not show accumulation into the BFA bodies (Fig 5f), suggesting that DUC1-GFP is not stably associated with early endosomes.

To examine the importance of the C2 domain for membrane-association of DUC1, we generated constructs of DUC1 expressing the full-length (FL), DUB domain, or C2 domain as RFP-fusion. When expressed in Arabidopsis root cell-derived protoplasts, 60% of the cells showed DUC1(FL)-RFP signals associated with the plasma membrane, with additional cytosolic and nuclear staining, whereas the DUB domain alone fused to RFP did not show plasma membrane localization (Fig 5g). In contrast, the C2 domain alone showed association with the plasma membrane in all analyzed cells (Fig 5g). These results shows that the DUC1 C2 domain is necessary and sufficient for the plasma membrane association of DUC1. Since the full-length DUC1 showed less efficient association with the plasma membrane, it can be speculated that the region outside of the C2 domain contribute to determining the localization of DUC1.

Other plasma membrane-localized DUBs OTU11 and OTU12 bind anionic lipids in membranes through their polybasic motif near the catalytic site. DUB activities of OTU11 and OTU12 are barely detectable in gel-based assays, whereas preincubation with liposomes leads to stimulation of their DUB activity ^12^. To test the possibility that DUC enzyme activity is also regulated by membrane binding, we compared the DUB activity of DUC1, DUC3, and OTU11. In contrast to OTU11, hydrolysis of di-UB to mono-UB were detectable for DUC1 and DUC3 (Supplementary Fig 3f), showing that recombinant DUCs are more active than OTU11. To test whether DUC activity is also affected by lipid binding, we conducted DUB assays with DUC3 preincubated with PI(4,5)P_2_-containing liposomes under the DUB:di-UB ratio typically used for OTU11 (Supplementary Fig 3g and h). The catalytic activity of DUC3 was not enhanced by the binding to PI(4,5)P_2_-containing liposomes, indicating that the DUB activity of DUC3 does not depend on its membrane binding.

### Role of DUC enzymes in endocytosis

To understand the physiological function of DUCs, we generated mutant lines. As the expression pattern suggested functional redundancy among the DUCs, we performed CRISPR/Cas9-based mutagenesis of *DUC1* in a T-DNA insertion mutant of *DUC3* and obtained two double mutant lines *duc1-1 duc3-1* and *duc1-2 duc3-1*. Both *duc1-1* and *duc1-2* have 1 bp insertions leading to a premature stop codon after 158 amino acids (Supplementary Fig 4a). The T-DNA insertion lies within an exon and causes the knock-out of *DUC3* (Supplementary Fig 4b to d). The primary root length of wild-type and *duc3-1* seedlings 5 days after germination was comparable. However, both double mutant lines *duc1-1 duc3-1* and *duc1-2 duc3-1* showed significantly longer primary root length compared to the wild type (Fig 6a to c), suggesting that DUC1 and DUC3 are involved in root growth regulation at the seedling stage. To analyse the effect of DUC1 and DUC3 knock-out on the global ubiquitylation profile, we compared the ubiquitylation profile of total extracts of wild-type, *duc1-1duc3-1*, *duc1-2duc3-1* and found that there are no apparent difference in the total ubiquitin amount (Supplementary Fig 4e), suggesting that loss of DUC1 and DUC3 does not affect the global ubiquitylation pattern but affect growth at the seedling stage.

**Figure 6:**
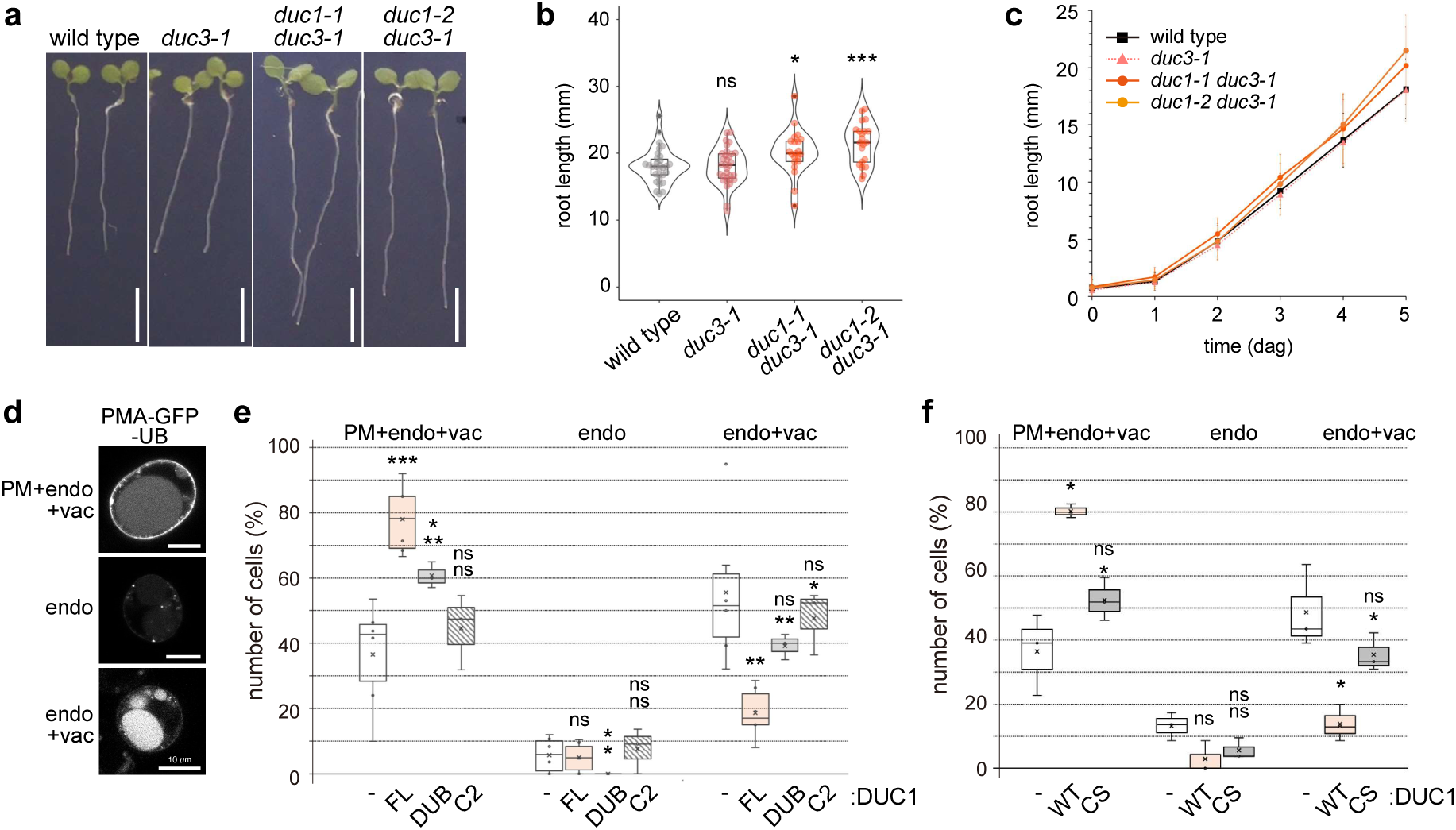
DUCs modulate plant growth and endosomal transport. **(a)** Representative photographs of wild-type, *duc1-1 duc3-1,* and *duc1-2 duc3-1* seedlings 5 days after germination (dag). Scale bars: 5 mm. **(b)** Primary root length of wild-type, *duc3-1, duc1-1duc3-1,* and *duc1-2duc3-1* seedlings 5 dag. Wild type/duc3-1 p=0.992 (ns: not significant), wild type/*duc1-1 duc3-1 p*=0.0244 (*: 0.01<*p*<0.05), wild type/*duc1-2 duc3-1 p*=0.000221 (***: *p*<0.001). The experiment was conducted three times, and one representative result is shown. **(c)** Root growth of wild-type, *duc3-1, duc1-1duc3-1,* and *duc1-2duc3-1* seedlings over time. The timepoint of germination was set to 0. **(d)(e)** DUC1 expression delays endosomal transport of the model cargo PMA-GFP-UB. Protoplasts expressing *PMA-GFP-UB* alone, *PMA-GFP-UB* with *35Spro:DUC1(FL)-RFP*, *35Spro:DUC1(DUB)-RFP*, or *35Spro:DUC1(C2)-RFP*, were analysed and cells were categorized according to the localization of PMA-GFP-UB as in (d). For PMA-GFP-UB and PMA-GFP-UB with *35Spro:DUC1(FL)-RFP*, six independent transformations were performed and for *35Spro:DUC1(DUB)-RFP,* and *35Spro:DUC1(C2)-RFP*, three experiments were performed. The results of all experiments are shown as a box plot. Center line, median; box limits, first and third quartiles; whiskers, 1.5x interquartile range; points, outliers. *p*-values for the PM+endosome+vacuole (PM+endo+vac) category: DUC1(FL)/control *p*=0.000629 (***: *p*<0.001), DUC1(DUB)/control *p*=0.0138 (*: 0.01<*p*<0.05), DUC1(DUB)/DUC1(FL) *p*=0.00957 (**: 0.001<*p*<0.01), DUC1(C2)/control *p*=0.431 (ns: 0.05<*p*), DUC1(C2)/DUC1(FL) *p*=0.0158 (*: 0.01<*p*<0.05); endo: DUC1(FL)/control *p*=0.784 (d: 0.05<*p*), DUC1(DUB)/control *p*=0.0449 (*: 0.01<*p*<0.05), DUC1(DUB)/DUC1(FL) *p*=0.0424 (*: 0.01<*p*<0.05), DUC1(C2)/control *p*=0.717 (ns: 0.05<*p*), DUC1(C2)/DUC1(FL) *p*=0.598 (ns: 0.05<*p*); endo+vac: DUC1(FL)/control *p*=0.00806 (**: 0.001<*p*<0.01), DUC1(DUB)/control *p*=0.135 (d: 0.05<*p*), DUC1(DUB)/DUC1(FL) *p*=0.0116 (*: 0.01<*p*<0.05), DUC1(C2)/control *p*=0.490 (ns: 0.05<*p*), DUC1(C2)/DUC1(FL) *p*=0.0176 (*: 0.01<*p*<0.05); two-tailed *t*-tests with no equal variance. **(e)** Protoplasts expressing *PMA-GFP-UB* alone, *PMA-GFP-UB* with 35Spro:DUC1(FL)-RFP, *35Spro:DUC1(DUB)-RFP*, or *35Spro:DUC1(C2)-RFP*, were analysed and cells were categorized according to the localization of PMA-GFP-UB as in (d). For *PMA-GFP-UB* and *PMA-GFP-UB* with *35Spro:DUC1(FL)-RFP*, six independent transformations were performed and for *35Spro:DUC1(DUB)-RFP* and *35Spro:DUC1(C2)-RFP,* three experiments were performed.

**Figure 7:**
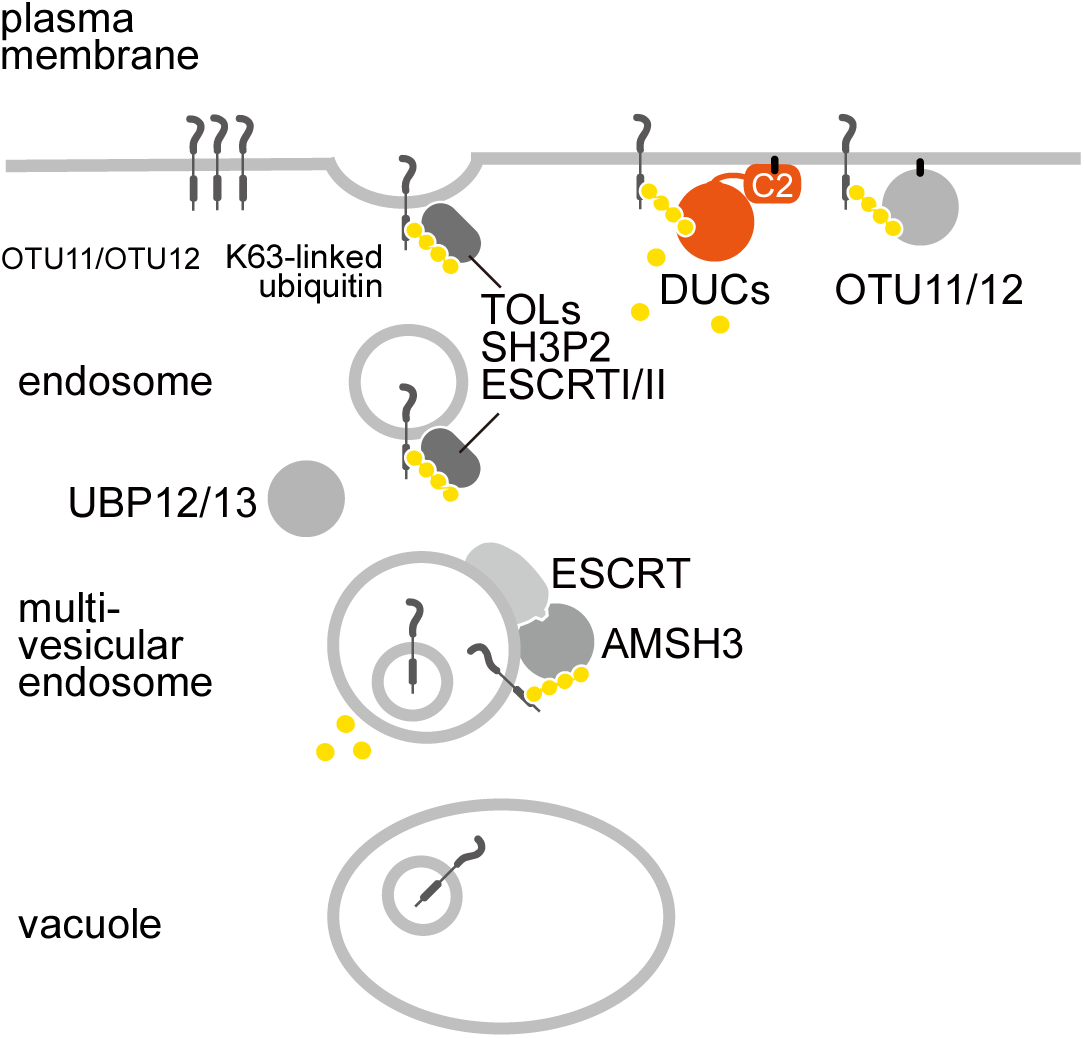
Model for DUC function. DUCs are plant-specific family of K63-specific DUBs. They localize to the plasma membrane by binding to phosphoinositides through their C2 domains. Overexpression of DUCs leads to the stabilization of a plasma membrane-localized endosomal model cargo in a DUB activity-dependent manner. This suggests that DUCs can modulate endosomal trafficking by removing the ubiquitin mark from endosomal cargoes, thereby preventing their recognition by endosomal ubiquitin readers such as the TOLs, SH3P2, or ubiquitin-binding ESCRT-I and ESCRT-II subunits, restoring them at the plasma membrane. The coordinated activities of endosomal transport-associated DUBs such as AMSH3, UBP12/UBP13, OTU11/OTU12 as well as the ubiquitination machinery and chain-specific readers control target degradation through the endosomal degradation pathway.

We next wanted to understand whether DUCs can affect endosomal trafficking in plants by modulating the ubiquitin mark which serves as a code for subsequent ESCRT-mediated endosomal degradation ^35^. We thus investigated whether the expression of DUC1 and DUC3 affect the endosomal transport of the endosomal model cargo PMA-GFP-UB ^36^. When PMA-GFP-UB is expressed on its own, it shows variable cellular localization profiles, which can be categorized into three patterns: Plasma membrane (+endosome/+vacuole), only on endosomes, on endosomes and the vacuole. In more than half of the cells, the GFP signals were found in the vacuole but not at the plasma membrane, indicating an efficient substrate transport from the plasma membrane to the vacuole via the endocytic machinery (Fig 6d and e). To test the effect of DUC1 and DUC3, we overexpressed DUC1 and DUC3 under a 35S promoter together with PMA-GFP-UB. Compared to the isolated PMA-GFP-UB expression, co-expression of DUC1 and DUC3 led to a significant increase of cells with plasma membrane signals, suggesting that DUC1 and DUC3 can impact the endosomal trafficking of PMA-GFP-UB (Fig 6e and Supplementary Fig 4f and g). This effect was less pronounced when only the DUB domain DUC1 was co-expressed and not observable when the C2 domain of DUC1 was co-expressed with PMA-GFP-UB (Fig 6e and Supplementary Fig 4g), suggesting that both the C2-domain mediated membrane localization and the DUB domain are important for the retention effect.

To corroborate that the enzymatic activity of DUC1 is important, we co-expressed wild type and a catalytic site cysteine to serine (CS) mutant of DUC1 with PMA-GFP-UB. Whereas the wild-type DUC1 overexpression led to the retention of PMA-GFP-UB at the plasma membrane, the CS variant did not affect the PMA-GFP-UB distribution significantly (Fig 6f and Supplementary Fig 4h). The CS mutation does not affect the cellular localization of DUC1, as both the wild-type and CS variants were observed at the plasma membrane to a similar extent (39.7% for wild type n=78, and 40.8 % for CS n=98, Supplementary Fig 4i). These results demonstrate that DUC1 can only interfere with the endosomal transport of a model cargo when its DUB activity is intact.

Altogether, our data show that DUCs are deubiquitinases localized to the plasma membrane in a C2 domain-dependent manner. Co-expression of DUC1 or DUC3 with a model endocytosis cargo leads to defects in endosomal transport to the vacuole, suggesting that DUC1 and DUC3 can modulate endosomal trafficking in Arabidopsis cells.

## Discussion

The discovery of the plant DUC family by an AlphaFold-based homology-agnostic screen demonstrates that new deubiquitinase families are still waiting to be discovered, even in well-studied model organisms such as *Arabidopsis thaliana*. The absence of detectable sequence similarity, even to the structurally related VTD family, suggests that the DUC enzymes could not have been identified by conventional homology-based screens ^22^. By contrast, the reaction with mono-ubiquitin-based probes (Fig 2a-c,g) suggests that probe-based pulldown screens can principally identify DUC members but might suffer from the generally low abundance and/or difficulty in detecting these proteins due to their membrane association.

The evolutionary relationship between the DUC family and established DUB classes is difficult to delineate. While performing a systematic comparison of all then-known DUBs and other cysteine protease families, we had noticed a distant sequence similarity between several of them, most prominently the USP, UCH, OTU, and Josephin families, and less significantly, the MINDY and ZUFSP families ^21^. This observation suggested the existence of a “proto-DUB” as a common ancestor of all cysteine DUBs and UBL proteases, but not of other papain-fold proteases. Only the VTD family, which at that time only comprised viral members, remained unconnected to other DUB classes in this analysis ^21^. After the expansion of the VTD family to members from fungi, protists, insects, and vertebrate transposons, it became clear that these enzymes share a common origin with the bacterial YopT family but differ in their active site architecture and their ubiquitin-directed specificity ^6^. The plant DUC family described here appears to be a third member of the YopT/VTD/DUC cluster. It shares with the VTD family a particularly similar papain-like fold and the propensity for linkage-selective deubiquitination, but has an active site that is much more YopT-like with the additional acquisition of an aromatic gatekeeper motif, which is found in most other DUB classes but absent from the non-ubiquitin cleaving YopT enzymes. VTD, DUC and YopT enzymes share considerable structural similarity but lack detectable sequence relationship and thus should be considered separate enzyme families.

Ubiquitin writers and readers are both detected at the plasma membrane in plant cells, suggesting that cargo modification and recognition can already start at the plasma membrane. For example, a number of K63-linkage assembling E3 ubiquitin ligases including the recently identified WAVY GROWTH proteins ^37^, RING DOMAIN LIGASEs (RGLGs) ^38^, IRT1 DEGRADATION FACTOR1 (IDF1) ^39^, member of the Plant-specific U-box proteins (PUBs) ^40^, were found to localize to the plasma membrane. Together with several other ubiquitin-binding proteins, TOM1-LIKEs (TOLs) take over the ESCRT-0 function in plants ^10^. Whereas ESCRT-0 in yeast and mammalian cells is found on early endosomes ^41,42^, the ubiquitin-binding SH3P2 and several TOL homologs are found at the plasma membrane ^43,44^. Thus, antagonizing ubiquitin eraser activity might already be required at the plasma membrane in plant cells.

While plasma membrane DUBs have not been reported in mammalian cells, plant OTU11/OTU12 are DUBs that localize to the plasma membrane through electrostatic interactions of their polybasic motif and anionic phospholipids ^12^. DUCs employ a C2 domain, a widespread membrane binding module, for plasma membrane localization. While OTU11/OTU12 activities are modulated by membrane binding, DUC activities do not depend on its membrane binding, and at least *in vitro*, DUCs are active without association to membranes.

DUCs are K63-specific DUBs, while OTU11/OTU12 also react with other linkages in addition to K63-linked ubiquitin chains. Given the dominance of K63 ubiquitin linkages in the endosomal protein trafficking pathway, the K63-linkage specificity of DUCs is indicative of their function in the regulation of K63-modified ubiquitin chains. DUCs affect the endosomal transport of a model cargo, suggesting that they can act on a broad range of plasma membrane cargos. The similarity of the assumed mode of functions between DUCs and OTU11/OTU12 may suggest functional redundancy between the families. Whereas OTU11 is expressed preferentially in the meristematic zone in roots, DUC1/DUC3 and OTU12 were found to be expressed in the elongation zone, implying a functional specialization between the DUBs. As OTU11/OTU12 and DUC1/DUC3 can affect the endosomal transport of the model cargo PMA-GFP-UB, and as neither protein family contains modules that could serve as further protein-protein interaction platforms, they are most likely able to act on a broad range of plasma membrane-resident target proteins.

## Materials & Methods

### Bioinformatics

Sequence alignments were generated using the L-INS-I method implemented in the MAFFT package ^45^. Generalized profiles were derived from multiple alignments and searched against the UniProt database (https://www.uniprot.org) using pfsearchV3 ^46^. HMM-to-HMM searches to establish sequence similarity between protein families were carried out using the HHSEARCH method ^47^. Structure predictions were performed using the ColabFold ^48^ implementation of Alphafold 2.3 ^23^. Structural database searches were performed using the DALI server ^30^. Structural superposition and calculation of RMSD values were performed using the TM-align server ^49^. Molecular modeling of individual protein complexes was performed using a local installation of Alphafold-3 ^33^, downloaded from GitHub (https://github.com/google-deepmind/alphafold3) using default databases and five seeds per run. Energy minimization was performed using a local installation of YASARA v25.1.13 with the YASARA force field ^50^.

### Cloning & Mutagenesis

AtDUC1-3 were amplified from *Arabidopsis thaliana* cDNA. All amplifications were performed by PCR using the Phusion High-Fidelity Kit (New England Biolabs). PCR fragments were cloned into the pOPIN-S vector ^51^ using the In-Fusion HD Cloning Kit (Takara Clontech). PpDUC1 was amplified from *Physcomitrium patens* cDNA (a gift from Ute Höcker and Maike Hansen, University of Cologne) and cloned accordingly. Point mutations were introduced using the QuikChange Lightning Kit (Agilent Technologies). A list of all primers used is provided in Supplementary Table 1. Constructs for ubiquitin-PA purification (pTXB1-ubiquitin^1–75^) were kindly provided by David Komander (WEHI, Melbourne). Human SUMO1^1–96^, SUMO3^1–91^, and ISG15^79–156^ were amplified by PCR with an N-terminal 3xFlag tag and cloned into the pTXB1 vector (New England Biolabs) by restriction cloning according to the manufacturer’s protocol.

Arabidopsis sequence information was retrieved from TAIR ^52^ (https://www.arabidopsis.org). The genomic sequences of the cloned genes were obtained by PCR from genomic DNA isolated from Arabidopsis seedlings. For all constructs of *DUC1* (At2g25460), the gene model *DUC1.2* was used, for all constructs of *DUC3* (At5g04860) the gene model *DUC3.1* was used.

To create C-terminally RFP-tagged transient expression constructs for cell culture-derived protoplasts pKV280 (*35Spro:DUC1-RFP*), pKV301 [*35Spro:DUC1*(*DUB*)-RFP], pKV307 [*35Spro:DUC1*(*C2*)-RFP], and pKV285 [*35Spro:DUC3-RFP*], full length *DUC1* was amplified with Primers KV481/482, the *DUC1* DUB domain with KV616/617, the *DUC1* C2-domain with KV567/568 and full length *DUC3* with KV484/486. The PCR-products were inserted with Gateway cloning into the entry vector pDONR 207 (Thermo Fisher Scientific) and subsequently into 35S-GW-RFP (MPI Cologne). For pTB308 (*DUC1 full length C318S*), the active site cysteine (C318) of DUC1 was mutated to serine by a Quickchange method using the Q5-Polymerase (NEB) and pKV280 as a template. To generate pTB270 [*NT*-DUC1-GST*], pTB273 [*GST-DUC3*], pTB234 [*GST-DUC1 (C2)*], pTB235 [*GST-DUC2 (C2)*], and pTB236 [*GST-DUC3 (C2)*], full length *DUC1* was amplified with TB580/TB581, full length *DUC3* with TB586/TB587, *DUC1 (C2)* was amplified with primers TB536/537, *DUC2 (C2)* with TB538/539, and *DUC3 (C2)* with TB540/TB541. Full length DUC3 was inserted in the SalI-site and the C2-domains were inserted in the BamHI/XhoI-sites of pGEX-6P1 (Thermo Fisher Scientific). Full length DUC1 was inserted via the NedI and SalI-sites into a pGEX-6P1 - derived vector which was modified for C-terminal tagging and contained an additional N-terminal NT*-tag ^53^. For recombinant expression, the coding sequence (CDS) of the Arabidopsis DUCs was synthesized by Eurofins and used as template.

The native promotor constructs, pKV311 (*DUC1pro:DUC1-GFP*) and pTB278 (*DUC3pro:DUC3-3xFlag-GFP*) were generated by Golden Gate cloning. The DUC1 promotor was amplified with primers KV532/533, the DUC3 promotor KV540/KV541, the CDS of DUC1 with KV534/KV535, and the genomic sequence of DUC3 with KV542/543. The BsaI-sites in the genomic sequence of DUC3 were mutated with primers TB589/TB590, KV546/KV547, KV548/KV549, and KV550/KV551 by overlap PCR. The promotors and genes were assembled with a NOS terminator, C-terminal GFP (DUC1), or C-terminal 3xFlag-GFP (DUC3) tag, a Basta resistance, and a *OLE1pro:OLE1-RFP* cassette into the pBB10 backbone vector ^54^. A list of new generated constructs is provided in Supplementary Table 2.

### Protein expression & purification

AtDUC1-3 and PpDUC1 were expressed using the pOPIN-S vector with an N-terminal 6His-SMT3-tag. *Escherichia coli* (Strain: Rosetta DE3 pLysS) were transformed with the respective constructs and 4-6□l cultures were grown in LB medium at 37□°C until the OD_600_ of 0.8 was reached. The cultures were cooled to 18□°C, and protein expression was induced by the addition of 0.1□mM isopropyl β-d-1-thiogalactopyranoside (IPTG). After 16□h, the cultures were harvested by centrifugation at 5000□×□*g* for 15□min.

After freeze-thaw, the pellets were resuspended in binding buffer (300□mM NaCl, 20□mM TRIS pH 7.5, 20□mM imidazole, 2□mM β-mercaptoethanol) containing DNase and lysozyme, and lysed by sonication using 10 s pulses at 50 W for a total time of 10 min. Lysates were clarified by centrifugation at 50,000□×□*g* for 1□h at 4□°C, and the supernatant was used for affinity purification on HisTrap FF columns (GE Healthcare) according to the manufacturer’s instructions. 6His-Smt3-tags were removed by incubation with SENP1^415-644^. The proteins were simultaneously dialyzed using the binding buffer. The liberated affinity tag and His-tagged SENP1 were removed by a second round of affinity purification using HisTrap FF columns (GE Healthcare). All proteins were purified using size exclusion chromatography (HiLoad 26/600 Superdex 75pg) in 20□mM TRIS pH 7.5, 150□mM NaCl, 2□mM dithiothreitol (DTT), concentrated using VIVASPIN 20 Columns (Sartorius), flash frozen in liquid nitrogen, and stored at −80□°C. Protein concentrations were determined using the absorption at 280 nm (A_280_) and the extinction coefficients of the proteins derived from their sequences.

### Enzymatic generation of activity-based probes

Ub^1-75^-PA, ISG15^79-154^-PA, SUMO1-PA, SUMO3-PA activity-based probes were expressed as C-terminal intein fusion proteins, as described previously^16^. In brief, intein fusion proteins were affinity-purified from clarified lysates in buffer A (20 mM HEPES, 50 mM sodium acetate pH 6.5, 75 mM NaCl) using Chitin Resin (New England Biolabs), following the manufacturer’s protocol. On-bead cleavage was performed by incubating with cleavage buffer (buffer A containing 100□mM MesNa (sodium 2-mercaptoethanesulfonate)) for 24□h at room temperature (RT). The resin was extensively washed with buffer A, and the pooled fractions were concentrated and subjected to size exclusion chromatography (HiLoad 16/600 Superdex 75pg) with buffer A. To synthesize the propargylated probe, 300□µM Ub/Ubl-MesNa was reacted with 600□µM propargylamine hydrochloride (Sigma Aldrich) in buffer A containing 150□mM NaOH for 3□h at RT. Unreacted propargylamine was removed by size-exclusion chromatography, and the probe was concentrated using VIVASPIN 20 columns (3□kDa cutoff, Sartorius), flash frozen, and stored at −80□°C. NEDD8-PA was a kind gift from David Pérez Berrocal and Monique Mulder (Department of Cell and Chemical Biology, Leiden University).

### Chain generation

Untagged Met1-linked diubiquitin was expressed as a linear fusion protein and purified using ion-exchange chromatography and size-exclusion chromatography. Wild-type 6His-tagged Met1-linked diubiquitin was expressed as linear fusion proteins and purified using HisTrap affinity purification and size exclusion chromatography. K11-, K48-, and K63-linked ubiquitin chains were enzymatically assembled using UBE2SΔC (K11), CDC34 (K48), and Ubc13/UBE2V1 (K63) as previously described ^55,56^. In brief, ubiquitin chains were generated by incubating 1□µM E1, 25□µM of the respective E2, and 2□mM ubiquitin in reaction buffer (10□mM ATP, 40□mM TRIS (pH 7.5), 10□mM MgCl_2_, 1□mM DTT) for 18□h at RT. The respective reactions were stopped by 20-fold dilution in 50□mM sodium acetate (pH 4.5), and chains of different lengths were separated by cation exchange using a Resource S column (GE Healthcare). Elution of different chain lengths was achieved using a gradient from 0 to 600□mM NaCl.

### Crystallization

AtDUC1^277-546^ was crystallized by sitting-drop vapor diffusion using commercially available sparse matrix screens. 96 well crystallization plates containing 30□µl of the respective screening conditions were mixed with 10□mg/ml protein in the ratios 1:2, 1:1, and 2:1 in 300□nl drops. Initial crystals appeared in PEG/Ion A1 (20 % v/v polyethylene glycol 3350, 0.2 M sodium fluoride) at 20°C and were cryoprotected in a 50 % saturated glucose in aforementioned crystallization solution.

### Data collection, phasing, model building, and refinement

Crystals were measured at the X06SA (PXI) beamline at the Swiss Light Source, Paul-Scherrer Institute Villigen, Switzerland. Diffraction data was processed using XDS^57^ and analyzed using pointless ^58^. Data showed strong signals with mean I/sigI > 2 at a resolution of 2.0 Å, for improved maps data up to a significant CC½ greater than 14.2% was retained^59^. Molecular replacement was performed using PHASER ^60^ with a predicted structure from the Chai-1 Server ^61^ as a search model. Initial models were rebuild using the modelcraft pipeline ^62^ and manually correct using iterative cycles of assisted model building with COOT ^63^ and refinement using either phenix.refine ^64^ or REFMAC ^65^. Data collection and refinement statistics are given in Supplementary Table 3.

### Activity-based probe assays

DUBs were prediluted to 2× concentration (10□µM) in reaction buffer (20□mM TRIS pH 7.5, 150□mM NaCl and 10□mM DTT) and combined 1:1 with 100□µM activity-based probes for 18 hours at 4 °C. The reaction was stopped by the addition of 2x Laemmli buffer, and analyzed by SDS-PAGE using Coomassie staining.

### Ubiquitin chain cleavage

DUBs were prediluted in 20□mM TRIS pH 7.5, 150□mM NaCl, and 10□mM DTT. The cleavage was performed at 20 °C for the indicated time points with 0.5 µM DUB and 25□µM diubiquitin (M1, K11, K48, K63 synthesized as described above, K6, K29, K33 purchased from Biomol, K27 from UbiQ. The reactions were stopped with 2x Laemmli buffer, resolved by SDS-PAGE and Coomassie stained.

For the comparison of DUCs and OTU11 in DUB assays, recombinant proteins were expressed in and purified from the *E.coli* Rosetta (DE3) strain (Merck) [*F-ompT hsdS* (r^-^ m^-^) *gal dcm (DE3) pRARE (Cam^R^)*]. GST-fused proteins were purified with ProtinoTM Glutathione Agarose 4B (Macherey-Nagel). 1 µM of GST-DUC3, DUC1-GST, and GST-OTU11 was each mixed 50 µM of K63-linked diubiquitin (UbiQ) in DUB assay buffer (20 mM Tris pH 7.5, 150 mM NaCl, 10 mM DTT) and incubated at 21°C. The reaction was stopped by adding 5 µL 5xSDS buffer at the indicated time points.

### Plant material and growth

*duc1-1* and *duc1-2* were generated using CRISPR/Cas9-mediated mutagenesis. Guide RNAs for *DUC1* and *DUC2* were inserted with primers KV591 and KV593, respectively, in the vector pHEE2E-TRI ^66^. The resulting construct *pKV333* was transformed in the *duc3-1* T-DNA insertion line. Transformants were selected with hygromycin and high-resolution melting analysis using the precision melt supermix (Biorad) and primers KV579/KV580. The mutation was confirmed by sequencing of the PCR product generated with primers KV658/KV659. The At5g04860 T-DNA insertion line *duc3-1* (WiscDsLox453-456A22, background Col-2) was obtained from NASC and genotyped with primers KV560/KV561 (wild type) and KV560/p745 (T-DNA insertion). For the RT-PCR, RNA was extracted from seedlings using the NucleoSpin RNA Plant kit (Macherey-Nagel) and reverse-transcribed with the M-MuLV reverse transcriptase (NEB). In the RT-PCR, the *DUC3* transcript was amplified with primers KV599/KV600, the control gene *ACTIN2* was amplified with primers *ACT2* fw/rv. Plant constructs were transformed by the Agrobacterium-mediated floral dip method. pKV291 and pKV311 were transformed in Col-0, pTB278 was transformed in Col-0 and *duc3-1*.

Arabidopsis seeds were surface sterilized in a solution of 1% NaOCl, stratified at 4 °C in dark for 1 to 3 days and grown under long day (16 hours light / 8 hours dark) at 21°C in ½ MS [2.15 g/L Murashige and Skoog medium including vitamin B5 (Duchefa), 0.5 g/L MES (Roth), pH 5.7].

For the root length measurement, *duc3-1* and progenies of a wild-type sibling obtained from a segregating population of *duc3-1* were grown for 7 days in long day on 1/2 MS after stratification for 3 days. The length of primary roots was measured with the SPIRO system ^67^ using the Spiro root growth macro in Fiji ^68^ and the Spiro R (https://www.r-project.org/) scripts. Graphs were prepared using Excel (Microsoft) or RStudio (http://www.rstudio.com/). Significance was tested using t-test (two-tailed, no equal variance) in Excel (Microsoft).

### Lipid overlay assay

Lipid overlay assays were performed using PIP Strips™ (Thermo Fisher Scientific) that were blocked for 5 to 7 hours at 4 °C in a PIP Strip buffer (20 mM Tris pH7.4, 150 mM NaCl, 0.02 % Tween, 3% fatty acid free bovine serum albumin, with 10 µM CaCl_2_ or 5 mM EGTA as indicated). After the blocking, the membranes were incubated with 1 µg/mL GST-fused proteins overnight at 4 °C. The lipid-bound proteins were detected using an anti-GST antibody dissolved in antibody buffer (20 mM Tris pH7.4, 150 mM NaCl, 0.02% Tween, 1% fatty acid free bovine serum albumin, 10 µM CaCl_2_).

### Liposome preparation and liposome sedimentation assay

Lipid vesicles were prepared as described previously ^69^. For the liposome sedimentation assay, freshly prepared liposomes [80% PC and 20% PE or 75% PC, 20% PE, and 5% PI3P or PI4,5P_2_ (lipids from Merck)] were mixed with 5 μg of recombinant protein and incubated for 15 min at room temperature in a liposome buffer [20 mM HEPES (pH7.4), 150 mM NaCl, 10 µM CaCl_2_, 50 mM sucrose]. For the sedimentation assay, the mixture was then centrifuged for 25 min in a Beckman TLA120.1 rotor at 100,000 ×*g* at 4°C. The supernatant and the pellet fraction were analysed by SDS-PAGE and TCE staining. The gels were imaged and quantified using Amersham Imager 600 (GE Healthcare).

For DUB assays with liposomes, 1.5 µM of GST-DUC3 in DUB assay buffer (20 mM Tris pH 7.5, 100 mM NaCl, 10 µM CaCl_2_, 2 mM DTT) with liposomes with and without PI(4,5)P_2_ in for 15 minutes at room temperature. After the incubation, 10 µL of the DUB-liposome mixture was mixed with 5 µL of 3 µM K63-linked di-ubiquitin (Boston Biochem) solution in DUB assay buffer (DUB:di-ubiquitin=1:1) and incubated at 21°C. The reaction was terminated by adding 5 µL of 5xSDS buffer at the indicated time points and the samples were analysed on NuPAGE 4-12% gradient gels (Thermo Fisher Scientific). The gels were stained with SYPRO-Ruby and the intensity of the ubiquitin bands was quantified with the Amersham^TM^ Imager 600 (Cytiva).

### Microscopy and image analyses

Protoplast transformations were done as described before ^43^. The confocal microscopy analysis of the protoplasts was done with the LSM700 with an 63x/1.40 PlanApochromat (oil) objective. The GFP-signal was excited with the 488 nm Diode-laser, the RFP-signal with the 555 nm Diode laser. The emission signal was collected with a 639 nm bandpass filter. Arabidopsis seedlings were imaged with the LSM880 with an 63x/1.40 PlanApochromat (oil) objective. For the excitation of GFP and RFP, the 488 nm Argon ion laser and the 561 nm diode-pumped solid-state laser were used, respectively. Images were obtained with the ZEN black software (Zeiss), and processed with Fiji ^68^, or OMERO figure ^70^. To stain the plasma membranes, 5- to 7-day-old seedlings were mounted in 2 µM FM4-64 in liquid ½ MS Seedlings and imaged directly. For the BFA treatment, seedlings were incubated with 50□µM brefeldin A (BFA, Sigma Aldrich) and 2 µM FM4-64 for 60□min at room temperature in the dark in liquid ½ MS.

## Material Availability

All biological materials used in this study will be made available upon request. T-DNA insertion lines of *duc3-1* can be obtained from Nottingham Arabidopsis Stock Centre (NASC) or Arabidopsis Biological Resource Center (ABRC).

## Data Availability

X-ray model coordinates and structure factors have been deposited at the PDB database under the accession code pdb_00009t71. Source data underlying the findings of this study are provided with this article.

## Acknowledgments

We thank Claudia Poschner and Justinian Bartz for expert technical assistance. This work was supported by the DFG (Grant HO 3783/3-2 to KH, 496470458 to EI). Mass spectrometry was performed at the Proteomics Facility of the CECAD, University of Cologne. Crystals were grown using equipment of the Cologne Crystallization facility (C_2_*f*), which is supported by DFG grant INST 216/949-1 FUGG. We thank Felix Groh for help in establishing the *duc* mutants, Johanna Gebauer, Sina Bergemann, and Catherine Dury (University of Konstanz) for earlier contribution to the project, the gardeners of the botanical garden of the University of Konstanz for taking care of the plants, and the Bioimaging Centre for the help in confocal microscopy. The synchrotron data were collected at the Swiss Light Source, Paul-Scherrer-Institute, Switzerland, at beamline X06SA. We thank the staff for their support.

## Author contributions

V.L. and M.S-W. performed the biochemical experiments. K.V. and T.B. performed lipid binding assays and plant experiments. T.H. performed the crystallization experiments. J.G. collected the X-ray data and solved the structure, and U.B. supervised the crystallography. K.H. conceived the project and contributed bioinformatical analyses. K.H. and E.I. jointly supervised the study. All authors contributed to the manuscript.

---

**Supplementary Figure 1:**
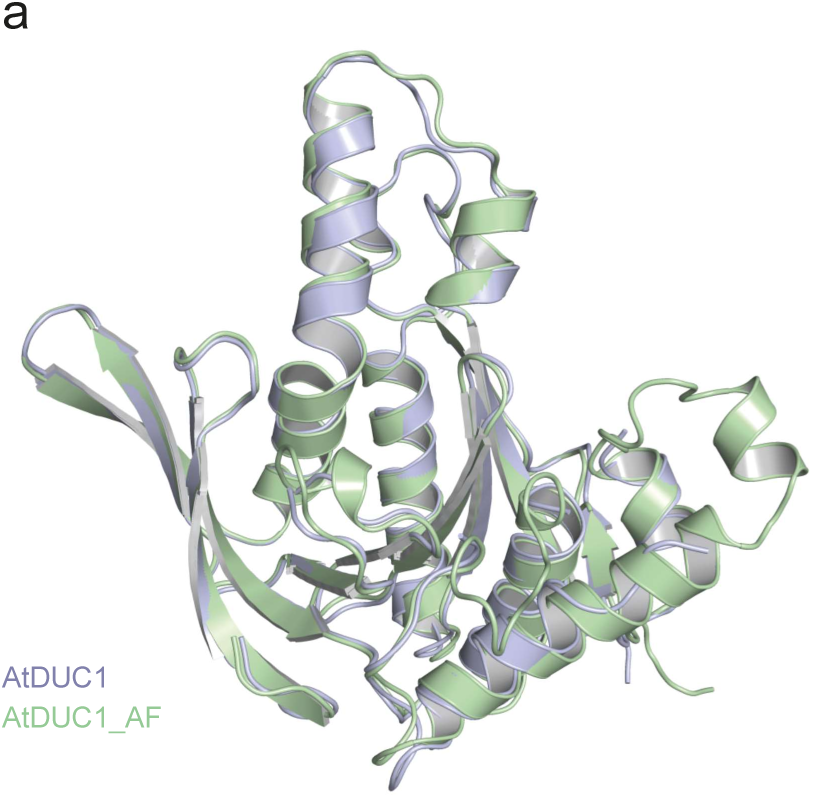
Comparison of the AtDUC1 structure with the AlphaFold model. a) The structures are shown in cartoon representation and coloured light purple (AtDUC1) and light green (AF model) respectively.

**Supplementary Figure 2:**
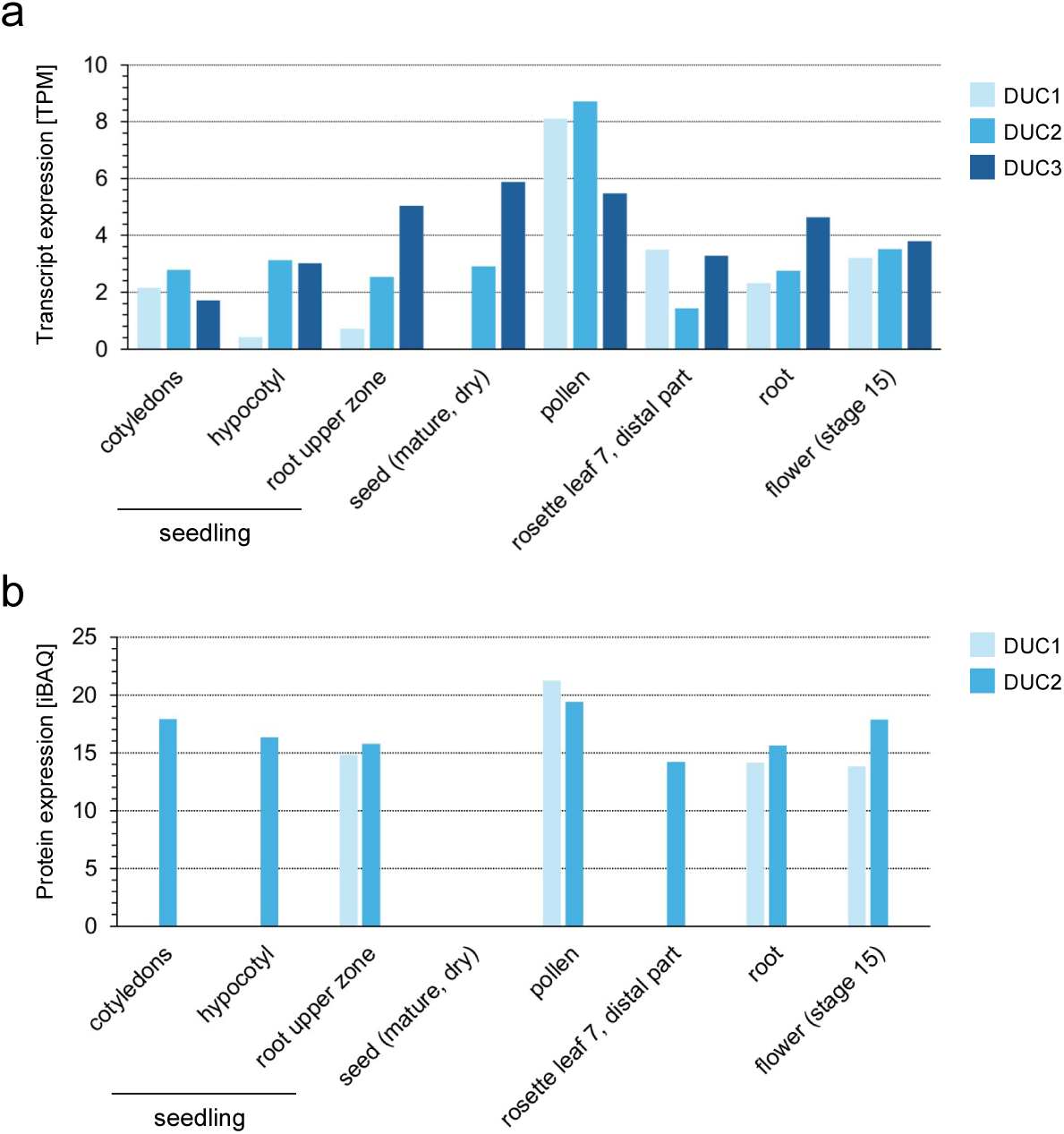
Expression of DUC1, DUC2, and DUC3. (a)(b) Transcript (a) and Protein (b) levels of DUC1, DUC2, and DUC3 in different Arabidopsis tissues (cotyledons, hypocotyl, root upper zone, seed, pollen, rosette leaf, root, and flower) as obtained from the ATHENA database (https://athena.proteomics.wzw.tum.de/master_arabidopsisshiny/). Note that DUC3 could not be detected in the proteomics dataset.

**Supplementary Figure 3:**
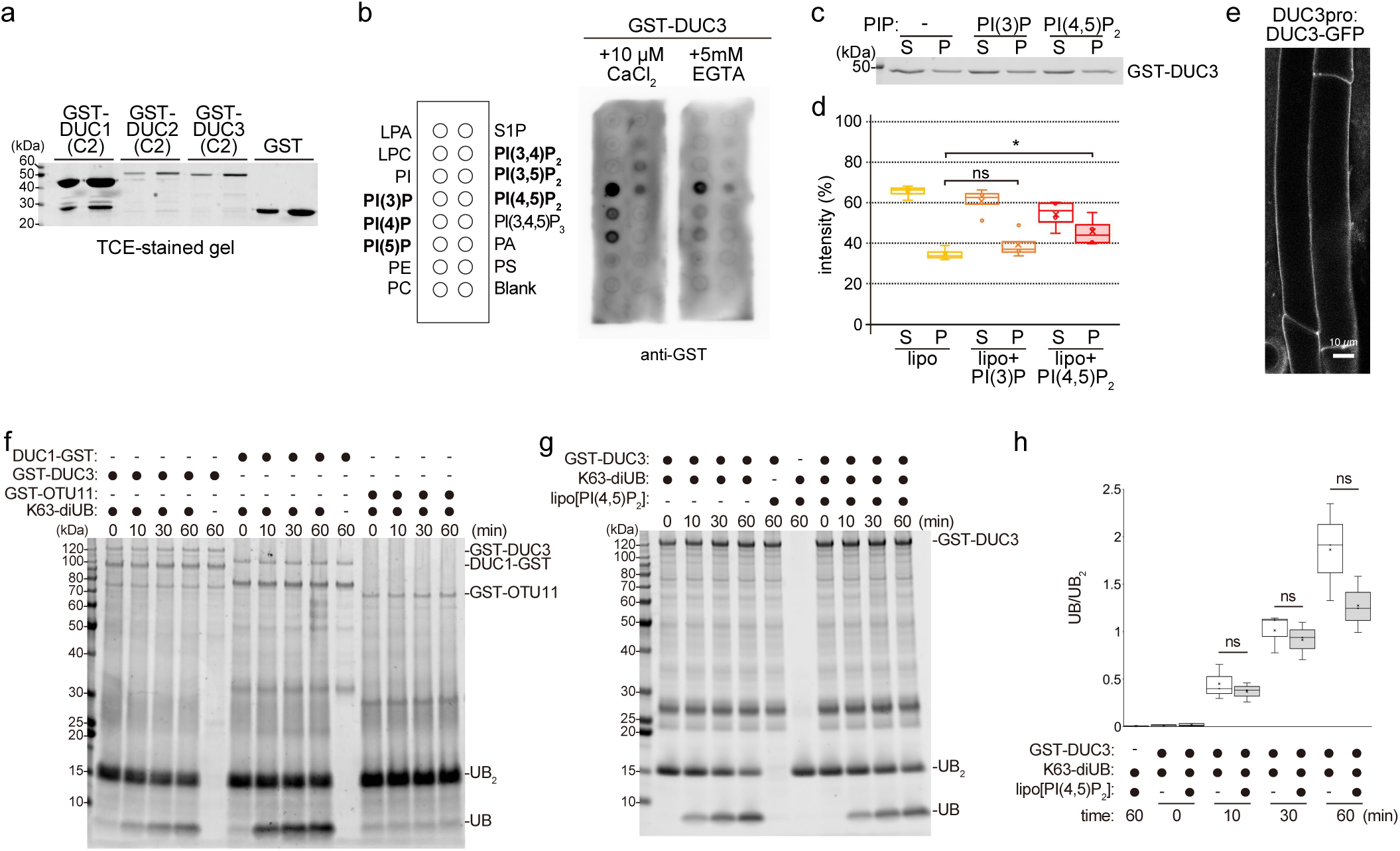
DUCT associate with lipid membrane. (a) Purification of GST and GST-fused DUC1, DUC2, DUC3. Purified proteins were subjected to SDS-PAGE and stained with TCE. (b) Lipid overlay assay of GST-fused GST-DUC3(C2) with either 10 µM CaCl2 or 5 mM EGTA. Lipid-bound proteins were detected with an anti-GST antibody. (c) Liposome sedimentation assay with GST-DUC3(C2). Recombinant GST-DUC3(C2) was incubated with liposomes (PC/PE) without PIPs or liposomes supplemented with 5% of PI(3)P or PI(4,5)P2 and centrifuged to obtain the supernatant (S) and pellet (P) fractions. SDS-PAGE gels were stained with TCE. Source data are provided as a Source Data file. The experiment was performed five times, and one representative result is shown. (d) Quantification of the result in (c). The signal intensity of the protein band in the supernatant-and pellet fractions was measured and the percentage in regard to the total combined signal intensity (S+P) was calculated. lipo: liposome. Box plot shows the results of quantification of five independent experiments. Center line, median; box limits, first and third quartiles; whiskers, 1.5x interquartile range; points, outliers. Comparison of the pellet fraction: lipo+PI(3)P/lipo p=0.264 (ns, not significant: 0.05<p), lipo+PI(4,5)P2/lipo p=0.0464 (*: 0.01<p<0.05). Two-tailed t-tests with no equal variance. Source data are provided as a Source Data file. (e) Image of DUC3pro:DUC3-GFP signals in the root epidermis cells of 5-day-old Arabidopsis seedling. Scale bars: 10 µm. (f) In vitro DUB assays using 1 µM of DUC1-GST, GST-DUC3, or GST-OTU11. K63-linked di-UB was added to the DUBs at a final concentration of 50 µM and incubated for the indicated time. The reactions were terminated by adding 5xSDS buffer. The experiment was conducted twice and one result is shown. (g) In vitro DUB assays using 1 µM of GST-DUC3 with or without preincubation with liposomes (PC/PE) containing 5% of PI(4, 5)P2. The DUBs were then incubated with 1 µM of K63-linked di-UB for the indicated time. The reactions were terminated by adding 5xSDS buffer and samples were analysed by SDS-PAGE. Gels were stained with TCE. The experiment was conducted three times, and one result is shown. (h) Quantification of the results as described in (g). The ration of mono ubiquitin (UB) and di-ubiquitin (UB2). Box plot shows the results of quantification of three independent experiments. Center line, median; box limits, first and third quartiles; whiskers, 1.5x interquartile range; points, outliers. Comparison of the pellet fraction: lipo+PI(4,5)P2/control, 0min p=0.592 (ns: 0.05<p); 30 min p=0.542 (ns: 0.05<p); 60 min p=0.177 (ns: 0.05<p). Two-tailed t-tests with no equal variance. Source data are provided as a Source Data file.

**Supplementary Figure 4:**
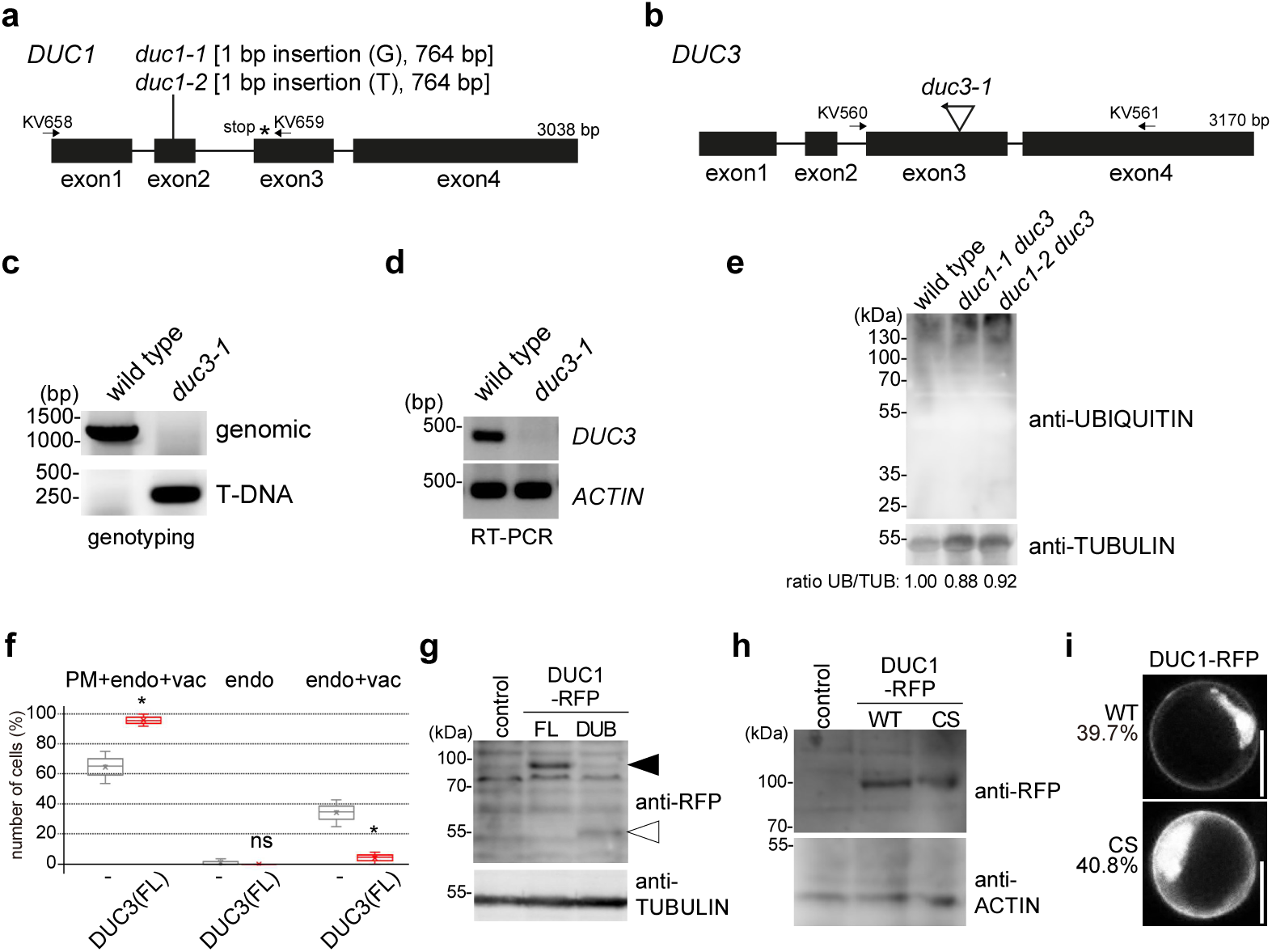
DUC1 and DUB3 modulate plant growth and endosomal transport. (a) Schematic representation of the DUC1 gene structure with the position of the duc1-1 and duc1-2 CRISPR events. The position of the premature stop codon in duc1-1 and duc1-2 is indicated with an asterisk. The positions of the genotyping primers are indicated with arrows. (b) Schematic representation of the DUC3 gene structure with the position of the duc3-1 T-DNA insertion. Arrow indicates the binding direction of the left border primer. The position of the genotyping primers is indicated with arrows. (c) Genotyping-PCR of wild-type and duc3-1 seedlings. PCR products for genomic fragments and T-DNA-containing fragments are shown. Source data are provided as a Source Data file. (d) RT-PCR of wild-type and duc3-1 seedlings. cDNA of wild-type and duc3-1 seedlings were analysed by PCR using gene-specific primers for DUC3 and ACTIN2 (control). Source data are provided as a Source Data file. (e) Total extracts of wild type, duc1-1, and duc1-2 were subjected to an immunoblot using an anti-ubiquitin antibody. An anti-TUBULIN antibody was used as loading control. The signal intensity of the anti-UBIQUITIN and anti-TUBULIN immunoblots were measured and the ratio for each lane are indicated below. (f) Protoplasts expressing PMA-GFP-UB alone or PMA-GFP-UB with 35Spro:DUC3(FL)-RFP were analysed and cells were categorized according to the localization of PMA-GFP-UB as in Figure 6a. For PMA-GFP-UB and PMA-GFP-UB with 35Spro:DUC3(FL)-RFP, three independent transformations were performed. The results of all experiments are shown as a box plot. Center line, median; box limits, first and third quartiles; whiskers, 1.5x interquartile range; points, outliers. p-values for the PM+endosome+vacuole (PM+endo+vac) category: DUC3(FL)/control p=0.0257 (*: 0.01<p<0.05); for the endosome (endo) category: DUC3(FL)/control p=0.423 (ns: 0.05<p); for the endo+vac category: DUC3(FL)/control p=0.0159 (*: 0.01<p<0.05); two-tailed t-tests with no equal variance. Source data are provided as a Source Data file. (g) Immunoblotting of total proteins extracted from cells used for analysis in Figure 6e. Total proteins were subjected to immunoblotting using an anti-RFP antibody. An anti-TUBULIN antibody was used as loading control on the same membrane. The C2 domain alone could not be detected. (h) Immunoblotting of total proteins extracted from cells used for analysis in Figure 6f. Total proteins were subjected to immunoblotting using an anti-RFP antibody. An anti-ACTIN antibody was used as loading control on the same membrane. (i) Representative images of cells expressing DUC1(WT)-RFP or DUC1(CS)-RFP at the plasma membrane. The percentage shows the number of cells with plasma membrane DUC1 signals. Scale bars: 10 µm.

