## Supplementary Tables for "DUCs are C2 domain containing plant-specific deubiquitinases stabilizing endocytic cargo at the plasma membrane"

**Supplementary Table 1: Primers used in this study**

| *primer name* | *sequence (5‘-3‘)* |
| --- | --- |
| DUC1 277 fw | GCGAACAGATCGGTGGTATCGAGGCAAACAAATGGGTAATGAAGGATTTAGTGAGCC |
| DUC1 546 rv | ATGGTCTAGAAAGCTTTAATCACTACTAGTCAATATAATATGATGCAAATCAATCTGAAGTCTCTCATGAAG |
| DUC1 C318A fw | CCGCCGGAGAAGCAGCAGCCGCGGCGGT |
| DUC1 C318A rv | ACCGCCGCGGCTGCTGCTTCTCCGGCGG |
| DUC1 H441A fw | AAATCTGCCTTTACCACGAAGAAAGCATCGTT CCAGCTAACAATATAAAC |
| DUC1 H441A rv | GTTTATATTGTTAGCTGGAACGATGCTTTCTTCGTGG TAAAGGCAGATTT |
| DUC1 D456A fw | CAATCTCTCCCCTAACGAAGCAATAACGCAAAAACCATC |
| DUC1 D456A rv | GATGGTTTTTGCGTTATTGCTTCGTTAGGGGAGAGATTG |
| DUC1 F442A fw | atattgttagctggaacgatcatgccttcgtggtaaaggcagatttag |
| DUC1 F442A rv | ctaaatctgcctttaccacgaaggcatgatcgttccagctaacaatat |
| DUC1 D373A fw | GAAACGATCGTTTCCAGAGCGAAATGCCTGTTTGGAA |
| DUC1 D373A rv | TTCCAAACAGGCATTTCGCTCTGGAAACGATCGTTTC |
| DUC1 E375A fw | TTGCGGAAACGATCGTTGCCAGATCGAAATGCCTG |
| DUC1 E375A rv | CAGGCATTTCGATCTGGCAACGATCGTTTCCGCAA |
| DUC1 D390A fw | ACCCGGTAAACGATTTAGCGGTACAAACCCTAACC |
| DUC1 D390A rv | GGTTAGGGTTTGTACCGCTAAATCGTTTACCGGGT |
| DUC1 T394R fw | GGGCTGAATAACCCCCTAAACGATTTATCGGTACAAACCC |
| DUC1 T394R rv | GGGTTTGTACCGATAAATCGTTTAGGGGGTTATTCAGCCC |
| DUC1 D536A fw | CTAGTCAATATAATATGATGCAAAGCAATCTGAAGTCTCTCATGAAGAA |
| DUC1 D536A rv | TTCTTCATGAGAGACTTCAGATTGCTTTGCATCATATTATATTGACTAG |
| DUC1 R290A fw | CAGCTTCGATTTTCCATCGGCGCTCACTAAATCCTT CATT |
| DUC1 R290A rv | AATGAAGGATTTAGTGAGCGCCGATG GAAAATCGAAGCTG |
| DUC1 R308A fw | CGGCTTGTTCGCTTGCTTGATCAATCGACGCTAAG TAAACTT |
| DUC1 R308A rv | AAGTTTACTTAGCGTCGATTGATCA AGCAAGCGAACAAGCCG |
| DUC1 E464A fw | AAGCTTGCTTGCATCCTGCAAACAATCTCTCCCCT |
| DUC1 E464A rv | AGGGGAGAGATTGTTTGCAGGATGCAAGCAAGCTT |
| DUC1 E513A fw | CTTTGCCGCCAACGCCGCCACCGGAAT |
| DUC1 E513A rv | ATTCCGGTGGCGGCGTTGGCGGCAAAG |
| DUC2 412 fw | GCGAACAGATCGGTGGTTTTGCGATTGGAAGTTGGGAGGAGAAAGAAGTGATAAGC |
| DUC2 702 rv | ATGGTCTAGAAAGCTTTATAGACTGGTAATCTCTGCGTTTGTGGTGTAATGAAACTCG |
| DUC3 427 fw | GCGAACAGATCGGTGGTGTTGTTGGTAGCTGGGAGACCAAAGAGATTATAAGCC |
| DUC3 733 rv | ATGGTCTAGAAAGCTTTAGTGGTGATGAAGGTGTTTCGTGTAGTGAAGCTCTATTTG |
| PpDUC 391 fw | GCGAACAGATCGGTGGTGGTGATGAAATTGACTGGCATCGACGGCAGG |
| PpDUC 748 rv | ATGGTCTAGAAAGCTTTAAACAGGAACCAACTCTGAGCAGGGCCAGG |
| KV481 DUC1 GW fw | GGGGACAAGTTTGTACAAAAAAGCAGGCTTGATGGCAAAGAAGGTGAGG |
| KV483 DUC1 C GW rv | GGGGACCACTTTGTACAAGAAAGCTGGGTCATCACTACTAGTCAATATAATATGATGC |
| KV484 DUC3 C GW fw | GGGGACAAGTTTGTACAAAAAAGCAGGCTTGATGGTGGTCAAAATGAAACAG |
| KV486 DUC3 C GW rv | GGGGACCACTTTGTACAAGAAAGCTGGGTCAACTTTAACCCCCAAAAGCT |
| KV532 DUC1pro GG A fw | ATATGGTCTCAGCGG CACCCATATCAAGTATTGGC |
| KV533 DUC1pro GG B rv | ATATGGTCTCACAGA AGATTTCAAAAGTTATTCCTCTG |
| KV534 DUC1 GG B fw | ATATGGTCTCATCTG TACA ATGGCAAAGAAGGTGAG |
| KV535 DUC1 GG C wo stop rv | ATATGGTCTCTGGTG CC ATCACTACTAGTCAATATAATATGATG |
| KV540 DUC3pro GG A fw | ATATGGTCTCAGCGG TCTCCAAAAATAAATAATTTAAAGC |
| KV541 DUC3pro GG B rv | ATATGGTCTCACAGA ACAATTCACAAGTGGCT |
| KV542 DUC3 GG B fw | ATATGGTCTCATCTG TACA ATGGTGGTCAAAATGAAACA |
| KV543 DUC3 GG C wo stop rv | ATATGGTCTCTGGTG CC AACTTTAACCCCCAAAAGC |
| KV546 DUC3 BsaI 168 m fw | GAGGAAGAGGCCTGTTGTT |
| KV547 DUC3 BsaI 168 m rv | AACAACAGGCCTCTTCCTC |
| KV548 DUC3 BsaI 873 m fw | TGTTCTATGGTCGCCTCTTTC |
| KV549 DUC3 BsaI 873 m rv | GAAAGAGGCGACCATAGAACA |
| KV550 DUC3 BsaI 1639 m fw | GTAGCTGGGAAACCAAAG |
| KV551 DUC3 BsaI 1639 m rv | CTTTGGTTTCCCAGCTAC |
| KV560 *duc3-1* GT fw | GTTCTGCTAATTGGGCTAGAGGC |
| KV561 *duc3-1* GT rv | CGGGTTGCTCAGACTTGTTCTT |
| KV567 DUC1 (C2) GW-C-fw | GGGGACAAGTTTGTACAAAAAAGCYAGGCTTGATGGCAAAGAAGGTGAGG |
| KV568 DUC1 (C2) GW-C-rv | GGGGACCACTTTGTACAAGAAAGCTGGGTCCTCAGAAAAAGTCACGTTGAC |
| KV579_DUC1_HRM1_fw | GATCGGAAAAGCATCGTTGGA |
| KV580_DUC1_HRM1_rv | CGAAAGTACGGAACCCTTTGA |
| KV591_DUC1_dcr_fw1 | ATATATGGTCTCGATTGTCGAAACAAGAATCGACGGGTTTTAGAGCTAGAAATAGC |
| KV593_DUC2_dcr_rv1 | ATTATTGGTCTCGAAACCGTATTTCCTCGTAACTAACAATCTCTTAGTCGACTCTAC |
| KV599 DUC3 RT fw | GCTTGGAGCAGGCGAAAG |
| KV600 DUC3 RT rv | CACATATTTCTCCACTCGGATG |
| KV614 DUC1 C318S fw | GCAGCATCCGCGGC |
| KV615 DUC1 C318S rv | GCCGCGGATGCTGC |
| KV616 DUC1 (DUB) GW C fw | GGGGACAAGTTTGTACAAAAAAGCAGGCTTGATCGAGGCAAACAAATGG |
| KV617 DUC1 (DUB) GW C rv | GGGGACCACTTTGTACAAGAAAGCTGGGTCATCACTACTAGTCAATATAATATGATG |
| KV658_DUC1_CrSeq_fw | ACGTGACGGTGAAGCC |
| KV659_DUC1_CrSeq_rv | CGGTTTCAACTACGGGG |
| TB536 DUC1 BamHI fw | ATATGGATCCATGGCAAAGAAGGTGAGGAAG |
| TB537 DUC1 C2 XhoI rv | TATACTCGAGTTAGTCGTCTGGTTCGGTTCTTAC |
| TB538 DUC2 BamHI fw | ATATGGATCCATGGTGGTGAAGATGATGAAGTG |
| TB539 DUC2-C2 XhoI rv | TATACTCGAGTTATGTTTCTGGAGTAGTTCTAAGCTC |
| TB540 DUC3 BamHI fw | ATATGGATCCATGGTGGTCAAAATGAAACAGATC |
| TB541 DUC3-C2 XhoI rv | TATACTCGAGTTACGATTCTTTAGGAGAAAACTGAAGGG |
| TB580 DUC1-Nde fw | ATATCATATGGAAAACCTGTATTTCCAATCGATGGCAAAGAAGGTGAGGAAGC |
| TB581 DUC1-Sal rv | ATATGTCGACATCACTACTAGTCAATATAATATGATGC |
| TB586 DUC3 SalI fw | ATATGTCGACTCATGGTGGTCAAAATGAAACAGATC |
| TB587 DUC3 SalI rv | ATATGTCGACTTAAACTTTAACCCCCAAAAGCTC |
| TB589 DUC3 Bsa mut fw | GGTAGCTGGGAGGCCAAAGAG |
| TB590 DUC3 Bsa mut rv | CTCTTTGGCCTCCCAGCTACC |
| ACT2 fw | CAAAGACCAGCTCTTCCATCG |
| ACT2 rv | CTGTGAACGATTCCTGGACCT |
| p745 | AACGTCCGCAATGTGTTATTAAGTTGTC |

**Supplementary Table 2: Plasmids generated in this study**

| *plasmid name* | | *description* | *vector backbone* | *source* |
| --- | --- | --- | --- | --- |
|  | | DUC1 277-546 WT | pOPINS | this study |
|  | | DUC1 277-546 C318A | pOPINS | this study |
|  | | DUC1 277-546 H441A | pOPINS | this study |
|  | | DUC1 277-546 D456A | pOPINS | this study |
|  | | DUC1 277-546 F442A | pOPINS | this study |
|  | | DUC1 277-546 D373A | pOPINS | this study |
|  | | DUC1 277-546 E375A | pOPINS | this study |
|  | | DUC1 277-546 D390A | pOPINS | this study |
|  | | DUC1 277-546 T394R | pOPINS | this study |
|  | | DUC1 277-546 D536A | pOPINS | this study |
|  | | DUC1 277-546 R290A | pOPINS | this study |
|  | | DUC1 277-546 R308A | pOPINS | this study |
|  | | DUC1 277-546 E464A | pOPINS | this study |
|  | | DUC1 277-546 E513A | pOPINS | this study |
|  | | DUC2 412-702 WT | pOPINS | this study |
|  | | DUC3 427-733 WT | pOPINS | this study |
|  | | PpDUC 391-749 WT | pOPINS | this study |
| pTB278 | | DUC3pro:DUC3-3xFlag-GFP | BB10 | this study |
| pKV311 | | DUC1pro:DUC1-GFP | BB10 | this study |
| pTB273 | | GST-DUC3 full length | pGEX-6P1 | this study |
| pTB234 | | GST-DUC1 C2-domain | pGEX-6P1 | this study |
| pTB235 | | GST-DUC2 C2-domain | pGEX-6P1 | this study |
| pTB236 | | GST-DUC3 C2-domain | pGEX-6P1 | this study |
| pKV301 | | 35SSpro:DUC1 (DUB) -RFP | 35S-GW-RFP | this study |
| pKV280 | | 35Spro:DUC1-RFP | 35S-GW-RFP | this study |
| pKV307 | | 35Spro:DUC1(C2)-RFP | 35S-GW-RFP | this study |
| pTB308 | | 35Spro:DUC1 C318S-RFP | 35S-GW-RFP | this study |
| pTB270 | | NT-(*TEV)-DUC1-GST | pGEX-6P1 | this study |
| pKV285 | | 35Spro:DUC3-RFP | 35S-GW-RFP | this study |
| pKV333 | | 2x CRISPR DUC1 gRNA1 and DUC2 gRNA2 | pHEE401E | this study |
| pKV119 | | GST-OTU11.2 | pGEX-6P1 | Ref ^12^ |
| PMA-GFP-UB | 35Spro:PMA-GFP-UB | |  | Ref ^36^ |
| pHEE2E-TRI | 2-single guide RNAs-EC1pro:zCas9 | |  | Ref ^66^ |

**Supplemental Table 3** Data collection and refinement statistics

|  | PVT3 |
| --- | --- |
| Data collection |  |
| Wavelength (Å) | 1.0000 |
| Space group | P 2_1_ 2_1_ 2_1_ (No. 19) |
| Cell dimensions |  |
| *a*, *b*, *c* (Å) | 54.89, 65.77, 76.84 |
| *α, β, γ* (°) | 90.00, 90.00, 90.00 |
| Resolution (Å) | 44.67 - 1.80 (1.87 - 1.80)^*^ |
| CC_½_ (%) | 99.9 (14.7)^*^ |
| *R*_merge_ (%) | 8.0 (165.0)^*^ |
| *R*_meas_ (%) | 9.1 (188.0) |
| Mean *I*/*I* | 12.93 (0.94) ^*^ |
| Completeness (%) | 97.5 (94.8) ^*^ |
| Multiplicity | 4.5 (4.3) ^*^ |
| Wilson B-factor | 27.89 |
| Refinement |  |
| No. reflections | 25749 (1292)^**^ |
| *R*_work/_ *R*_free_ (%) | 20.6 / 23.0 |
| No. atoms (non-hydrogen) |  |
| Protein | 1896 |
| Water | 61 |
| B-factors |  |
| Protein | 47.9 |
| Water | 41.1 |
| R.m.s deviations |  |
| Bond lengths (Å) | 0.05 |
| Bond angles (º) | 0.73 |
| Ramachandran |  |
| Favoured (%) | 96.6 |
| Allowed (%) | 3.4 |
| Forbidden (%) | 0 |

*Highest resolution shell is shown in parenthesis.

** Reflections used in R_free_
